# Cellular expression and functional roles of all 26 neurotransmitter GPCRs in the *C. elegans* egg-laying circuit

**DOI:** 10.1101/2020.04.23.037242

**Authors:** Robert W. Fernandez, Kimberly Wei, Erin Y. Wang, Deimante Mikalauskaite, Andrew Olson, Judy Pepper, Nakeirah Christie, Seongseop Kim, Michael R. Koelle

## Abstract

Maps of the synapses made and neurotransmitters released by all neurons in model systems such as *C. elegans* have left still unresolved how neural circuits integrate and respond to neurotransmitter signals. Using the egg-laying circuit of *C. elegans* as a model, we mapped which cells express each of the 26 neurotransmitter G protein coupled receptors (GPCRs) of this organism and also genetically analyzed the functions of all 26 GPCRs. We found that individual neurons express many distinct receptors, epithelial cells often express neurotransmitter receptors, and receptors are often positioned to receive extrasynaptic signals. The egg-laying circuit appears to use redundancy and compensation to achieve functional robustness, as receptor knockouts reveal few defects; however, increasing receptor signaling through overexpression more efficiently reveals receptor functions. This map of neurotransmitter GPCR expression and function in the egg-laying circuit provides a model for understanding GPCR signaling in other neural circuits.

## Introduction

Analysis of small neural circuits, such as those of genetically-tractable model organisms, has the potential to uncover principles that generalize to circuits of more complex nervous systems (Bargmann and Marder, 2013). Deeply understanding a neural circuit might involve mapping the synaptic connections between all its cells, identifying the neurotransmitters released by each neuron, identifying the neurotransmitter receptors present on each cell, and defining how signaling through each receptor affects the circuit’s function. This depth of understanding has not yet been achieved for any circuit.

A major challenge is to understand how neurotransmitters control neural circuit function by acting as neuromodulators through G protein coupled receptors (GPCRs) (Marder, 2012). Each neurotransmitter has multiple GPCRs: serotonin, for example, has 12 different GPCRs expressed in human brain that can couple to different G proteins to produce different effects (McCorvy & Roth, 2015). Neurotransmitters bind GPCRs with high affinity, allowing response to neurotransmitters released at a distance, an effect known as extrasynaptic or volume transmission (Fuxe et al., 2005; Agnati et al., 2006; Fuxe et al., 2012). Thus, mapping the synaptic connections between the neurons in a circuit is insufficient to understand the GPCR signaling that may occur between them.

*C. elegans* provides an opportunity to define how neural circuits use neurotransmitter GPCR signaling because: 1) A complete neural connectome delineates ∼7,000 synaptic connection between all 302 neurons in this organism (Albertson & Thomas, 1976; White et al., 1986; Xu et al., 2013); 2) A complete neurotransmitter map shows which of seven small-molecule neurotransmitters is released by each neuron (Serrano-Saiz et al., 2013; Pereira et al., 2016; Gendrel et al., 2016); 3) *C. elegans* has ∼26 homologs of human neurotransmitter GPCRs, 21 of which have been assigned as receptors for one of the seven neurotransmitters (while the remainder are “orphan” receptors without assigned ligands); and 4) Gene knockouts for all neurotransmitter GPCRs are available (Koelle 2018).

In this study, we analyzed the expression and functions of all 26 neurotransmitter GPCRs in the neural circuit that controls egg laying in *C. elegans*. This circuit serves as a model for neural G protein signaling since mutations that alter signaling by the neural G proteins Gα_o_, Gα_q_, and Gα_s_ strongly affect the frequency of egg laying (Schafer, 2006; Koelle, 2018). Also, the neurotransmitters serotonin, tyramine, octopamine, and dopamine, which act through GPCRs, have all been shown to functionally affect egg laying (Horvitz et al., 1986; Schafer and Kenyon, 1995; Hapiak et al., 2009; Vidal-Gadea et al., 2012; Collins et al., 2016; Nagashima et al., 2016). However, it remains to be determined how specific receptors for these neurotransmitters alter activity of the egg-laying circuit. Our analysis reveals features of neurotransmitter signaling that may generalize beyond the egg-laying circuit, and provides a path toward understanding how neurotransmitter signaling within a model neural circuit generate its dynamic pattern of activity to control a behavior.

## Results and Discussion

### GFP Reporter Strains to Identify Cells Expressing each Neurotransmitter GPCR

Our analysis includes 21 neurotransmitter GPCRs for monoamines and classical neurotransmitters, plus five “orphan” GPCRs (Figure 1A). Expression patterns for thirteen of these have been previously analyzed (Figure S4; Cho et al., 2000; Lee et al., 2000; Tsalik et al., 2003; Chase et al., 2004; Rex et al., 2004; Carnell et al., 2005; Dempsey et al., 2005; Carre-Pierrat et al., 2006; Suo et al., 2006; Wragg et al., 2007; Hapiak et al., 2009; Plummer, 2011; Gürel et al., 2012; Yemini et al., 2019) using GPCR gene promoter fragments of a few kilobases to drive green fluorescent protein (GFP) expression. However, recent work shows that obtaining accurate expression patterns often requires larger GFP reporter transgenes (Serrano-Saiz et al., 2013; Pereira et al., 2016; Gendrel et al., 2016) that include the full gene of interest and up to ∼20 kb of flanking genomic DNA on either side (Tursun et al., 2009).

**Figure 1:**
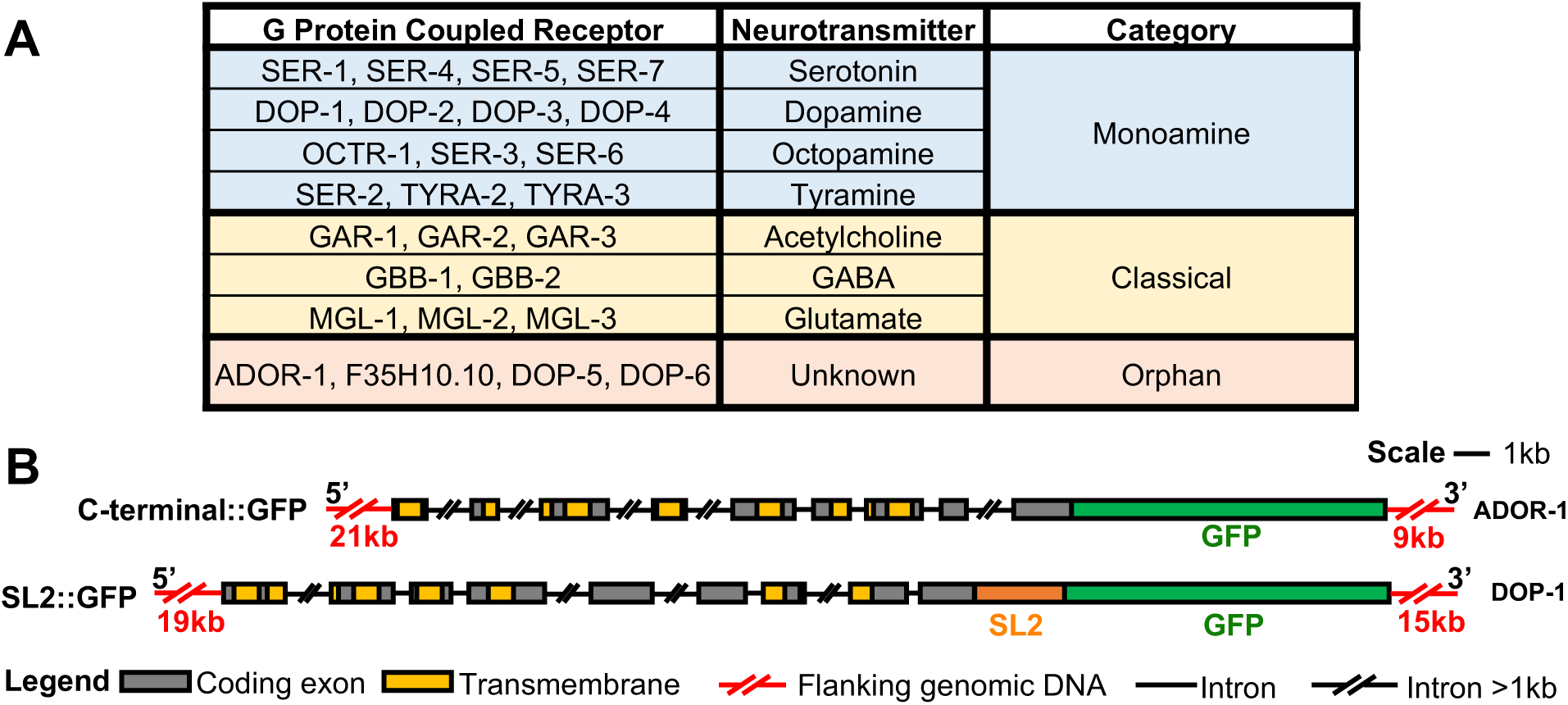
**GFP Reporter Transgenes for 26 Neurotransmitter GPCRs** (A) Small-molecule neurotransmitter GPCRs in *C. elegans*. (B) Schematics of GFP reporters used in this study. C-terminal::GFP constructs are illustrated by the reporter for the orphan receptor ADOR-1. The construct provides 21 kilobase (kb) of promoter region DNA 5’ to the coding exons and 9 kb of 3’ DNA. GFP coding sequences were inserted into the last ADOR-1 coding exon so that the transgene expresses the receptor with GFP fused to its C-terminus. SL2::GFP constructs are illustrated by the dopamine receptor DOP-1 reporter. This reporter is similar to that for ADOR-1 except that an SL2 trans-splicing signal followed by GFP coding sequences were inserted immediately downstream of the stop codon so that this reporter co-expresses the receptor and GFP as separate proteins. Analogous schematics of all 26 receptor::GFP reporter transgenes are found in Figure S1 and described in Table S1.

Therefore, we sought to generate large GFP reporter transgenes for the 26 neurotransmitter GPCR genes (Figures 1 and S1 and Table S1). Thirteen were already constructed (Sarov et al. 2012), and we generated seven more in a similar fashion. These “C-terminal::GFP” reporters express a GPCR with GFP fused at or very close to the C-terminus of the receptor (Figure 1B). For a few receptors, these C-terminal::GFP fusion proteins were strongly localized to neural processes and excluded from the cell bodies, making it difficult to identify the GFP-expressing neurons. So, for four such receptors (DOP-1, DOP-4, GAR-2, and F35H10.10) we generated “SL2::GFP” reporters (Figure 1B) which produce a single primary RNA transcript that is trans-spliced to produce separate GPCR and GFP mRNAs, so that the GFP produced fills out the cells that express the GPCR gene (Tursun et al. 2009). For the remaining two receptors (MGL-2 and SER-4), technical issues precluded use of large fosmid-based reporters, so we analyzed expression of smaller promoter::GFP reporters that had been previously generated (Tsalik et al., 2003; Gürel et al., 2012; Yemini et al., 2019).

The 26 GPCR::GFP reporter transgenes were each transformed into *C. elegans* to produce high-copy, chromosomally-integrated transgenes. Such transgenes are expressed in the same cells as the corresponding endogenous genes (Serrano-Saiz et al., 2013; Pereira et al., 2016; Gendrel et al., 2016), but at high enough levels that the GFP fluorescence produced is sufficient to identify even cells that normally express a GPCR gene at low levels.

### Cells of the Egg-Laying System

We first examined animals carrying transgenes that express mCherry in specific cells of the egg-laying system to generate accurate diagrams of its anatomy (Figure 2).

**Figure 2:**
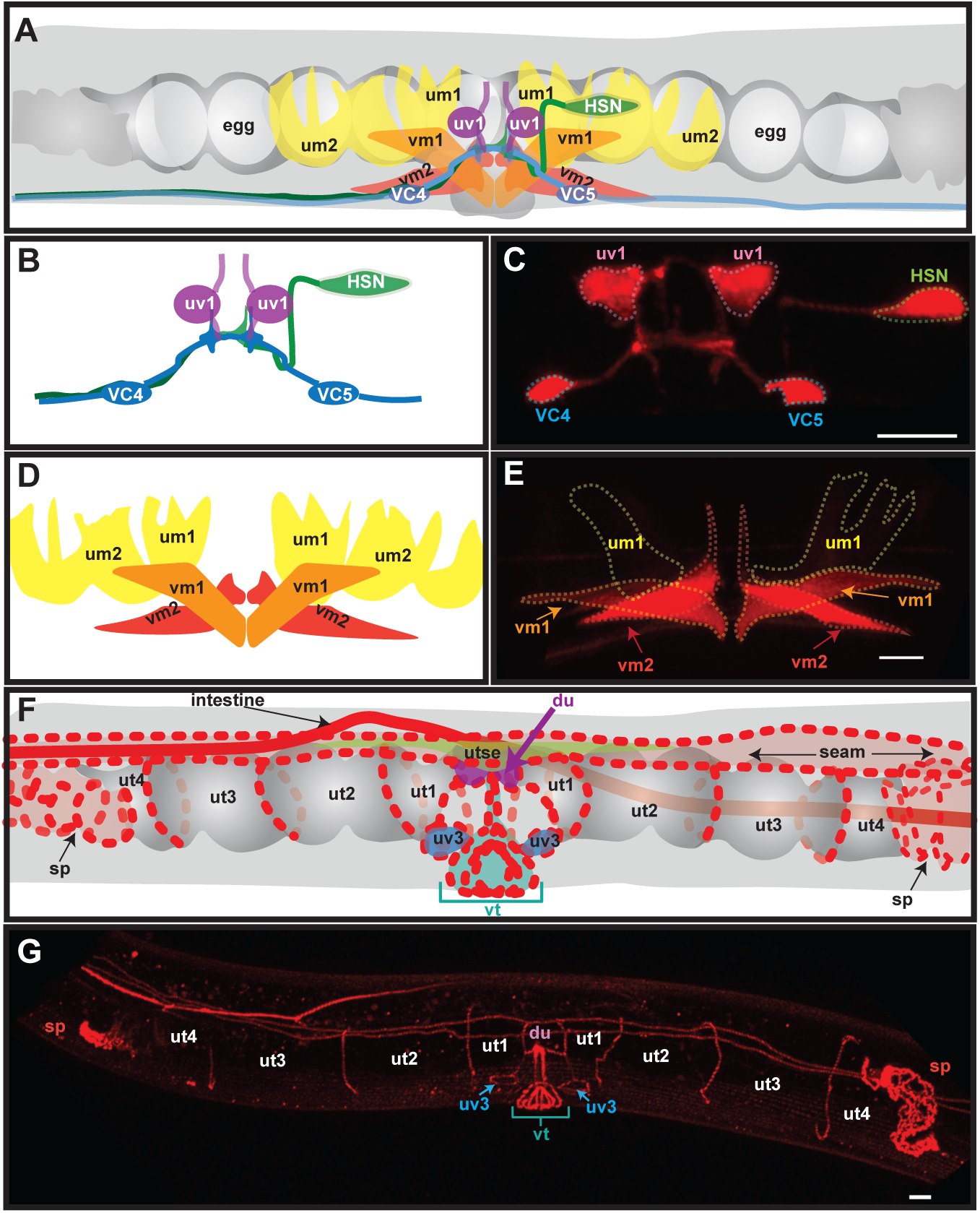
**Anatomy of the Egg-Laying System Visualized with mCherry Markers** (A) Schematic of the *C. elegans* egg-laying system. Neuron and muscle cells (colored and labeled with cell names) are overlaid on the uterus (grey) containing eggs (white). VC4 and VC5 neurons are at the ventral midline. All other cells depicted are on the left side of the animal and their equivalents on the right side of the bilaterally symmetric anatomy are not shown. (B) Schematic of the egg-laying neurons. (C) Confocal image of the egg-laying neurons labeled by the *ida-1::mCherry* marker. Dashed lines outline the cell bodies. (D) Schematic of the egg-laying muscles. (E) Confocal image of the egg-laying muscles labeled by the *unc-103e::mCherry* marker. um1 (outlined) and um2 (not indicated) are rarely and faintly labeled by *unc-103e::mCherry*, but even so labeling of vm1/vm2 cells provides landmarks to help identify um1 and um2. (F) Schematic of cells other than neurons and muscles of the egg-laying system. Red dashed or solid lines indicate cell junctions labeled by the *ajm-1::mCherry* marker. Besides cells of the egg-laying system, the *ajm-1::mCherry* marker also labels junctions of seam and intestinal cells. (G) Confocal image of the egg-laying system labeled by the *ajm-1::mCherry* marker. The following conventions are used in this and other figures in this work: Scale bars are 10 µm. Anterior is left and ventral down. Abbreviations used are du: dorsal uterine cell; HSN: hermaphrodite specific neuron; sp: spermatheca; um: uterine muscle cell; ut: uterine toroid cell; utse: uterine seam cell; uv: uterine ventral cell; VC: ventral cord type C neuron; vm: vulval muscle cell; vt: vulval toroid cells.

Neurons of the egg-laying system were labeled by *ida-1::mCherry* (Figures 2B and 2C). These neurons are: 1) The HSNs, which promote egg laying by releasing serotonin and neuropeptides (Waggoner et al., 1998; Shyn et al., 2003; Hapiak et al., 2009; Emtage et al., 2012; Brewer et al., 2019); 2) the uv1s, which inhibit egg laying by releasing tyramine and neuropeptides (Alkema et al., 2005; Collins et al., 2016; Banerjee et al., 2017); and 3) VC4/5, which release acetylcholine onto the vm2 egg-laying muscles (Duerr et al., 2001; Collins et al., 2016).

Muscle cells of the egg-laying system were labeled by *unc-103e::mCherry*, which is expressed in the vm1 and vm2 vulval muscle cells and was used as a landmark to identify the um1 and um2 uterine muscle cells (Figures 2D and 2E). Eggs are laid when vm1 and vm2 open the vulva and um1 and um2 muscle cells squeeze eggs out of the uterus through the vulva (Kim et al., 2001; Waggoner et al., 2001; Bany et al., 2003; Ringstad and Horvitz., 2008; Collins et al., 2016). There are also body wall muscles cells surrounding the egg-laying system (not shown in Figure 2 but referred to as bwm in later figures).

Other cells of the egg-laying system were identified using *ajm-1::mCherry*, which labels the junctions between cells of the uterus and vulva (Figures 2F-2H). Eggs are fertilized in the spermatheca (sp), and enter the uterus via the spermatheca-uterine (sp-ut) valve. The uterus itself comprises the ut1-ut4 uterine toroid cells and du cells that together form a tube to hold eggs. Eggs pass from the uterus into the vulva through a junction comprising the uterine seam (utse) and uterine-ventral (uv1 and uv3) cells, and the vulva itself comprises several types of vulval toroid (vt) cells (Schindler and Sherwood, 2014; Ghosh and Sternberg, 2014; Ecsedi et al., 2015).

### Analyzing GFP Expression by Neurotransmitter GPCR::GFP Transgenes

We sought to 1) accurately identify every cell of the egg-laying system that expresses any of the 26 neurotransmitter GPCR::GFP transgenes; 2) qualitatively record the GFP expression levels for each reporter in each cell, which may reflect expression levels of the endogenous receptors; and 3) record animal-to-animal variability in GFP expression levels from these reporters, which may reflect variability in expression of the endogenous receptors. Thus, we crossed each of the 26 neurotransmitter GPCR::GFP transgenes into the mCherry marker strains and collected three-dimensional confocal images for at least 10 young adult animals per double-labeled strain. In total, images of 809 animals were analyzed.

The process for identifying GFP-expressing cells and scoring GFP levels is illustrated for the octopamine receptor OCTR-1 in Video S1 and Figure 3. Each image was analyzed using 3D rotations, cuts through the image volumes, and mCherry markers to identify internal features. The strongest GFP expressing cells in a strain were scored as “+++”, those with weaker but easily detectable GFP as “++”, and those with GFP just above background as “+”.

**Figure 3:**
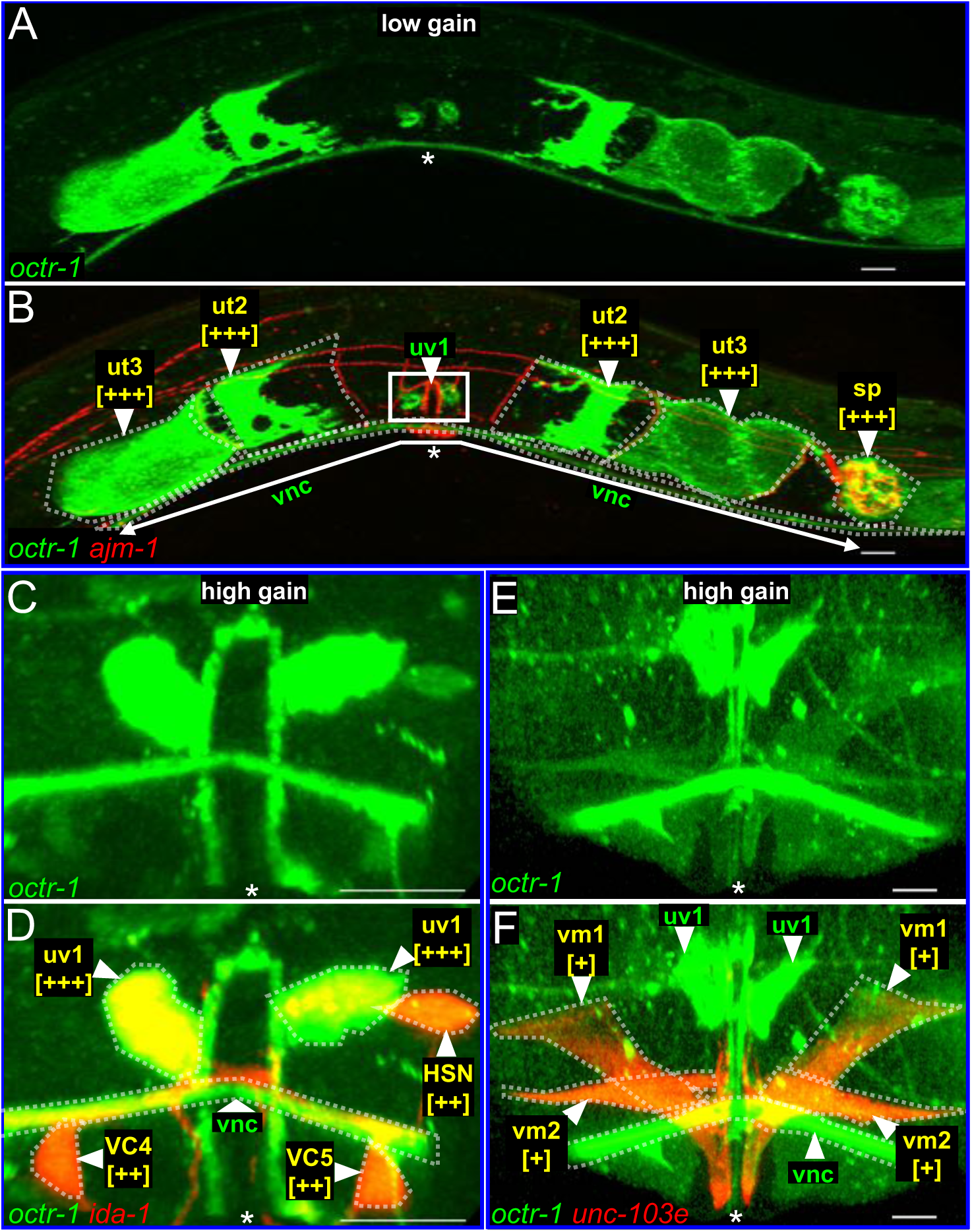
**Scoring Expression of a GFP Reporter for the Octopamine Receptor OCTR-1 in Cells of the Egg-Laying System** (A and B) Two-dimensional renderings of a confocal image of the egg-laying system in an *octr-1::gfp; ajm-1::mCherry* young adult showing GFP (A) or GFP and mCherry fluorescence (B). The ut2 uterine toroid cells are outlined by the *ajm-1::mCherry* marker, allowing them to be identified as GFP positive despite not being uniformly filled out by OCTR-1::GFP. The solid with arrowheads labels the ventral nerve cord (vnc), a bundle of axons that run past the vulva but that are not known to affect egg laying. In these and other figure panels, white asterisks indicate position of the vulval opening, cells labeled with both GFP and mCherry are named with yellow text, cells labeled with GFP only are named with green text, the number of “+” symbols indicates the relative intensity of GFP labeling, and identified cells are outlined by faint dotted lines. White box in (B) indicates the region shown at higher magnification in (C-F). (C and D) An *octr-1::gfp; ida-1::mCherry* animal with GFP fluorescence displayed at high gain to make ++ GFP labeling visible in the HSN and VC4/5 neurons. These neurons are double-labeled by *ida-1::mCherry* in (D). (E and F) An *octr-1::gfp; unc-103e::mCherry* animal with GFP fluorescence displayed at high gain and at a different focal plane than in (C and D) to visualize + GFP labeling in vm1 and vm2 egg-laying muscles.

For reporters that expressed a receptor with GFP fused to its C-terminus, the fusion proteins were sometimes subcellularly localized. An example is seen in Figure 3B, where OCTR-1::GFP clusters within the ut2 uterine toroid cells. It is unclear if such subcellular localization might reflect a similar localization of the corresponding endogenous receptors; thus, we chose not to interpret such subcellular localization in this work.

### Neurotransmitter GPCR Expression in the *C. elegans* Egg-Laying System

Figure 4 shows two-dimensional image renderings illustrating a subset of findings, while Figure S3 shows a larger set of images for each of the 26 neurotransmitter receptors. Figure 5 summarizes all the findings from all 809 images analyzed. In total, our results show 139 cases in which a neurotransmitter GPCR is expressed in a specific cell type of the egg-laying system.

**Figure 4:**
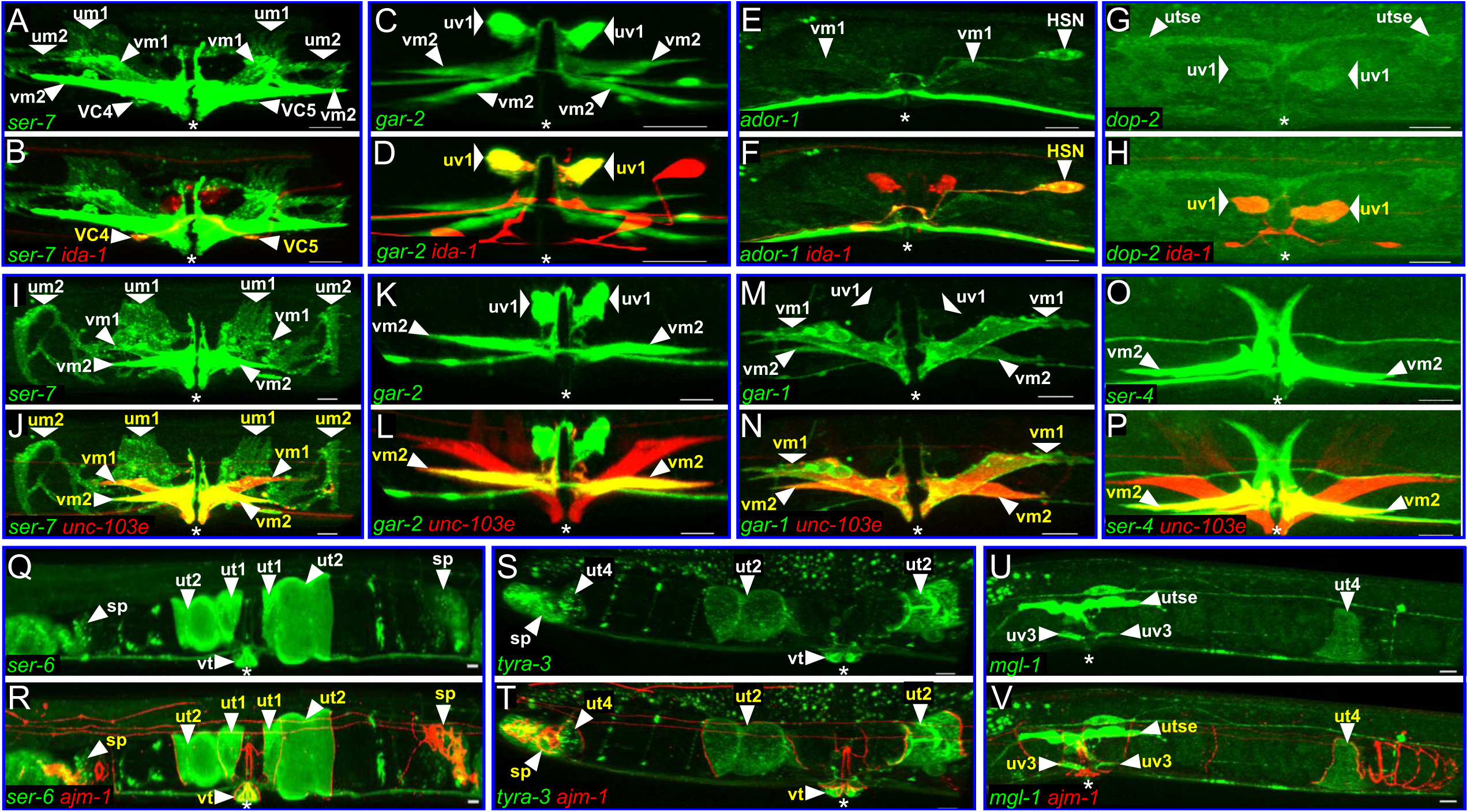
**Examples of Neurotransmitter GPCR::GFP Expression in Neuronal, Muscle, and Other Cells of the Egg-Laying System** (A-H) Examples of GPCR::GFP expression in neurons of the egg-laying system. Upper panel of each image pair shows reporter GFP reporter fluorescence for the receptor named in green, with all GFP-labeled cells of the egg-laying system indicated by white cell names and arrowheads. Lower panels show GFP plus the *ida-1::mCherry* fluorescence used to confirm identity of egg-laying neurons, with only the double-labeled neurons indicated by yellow names and white arrowheads. (I-P) Examples of GPCR::GFP expression in muscle cells of the egg-laying system. Images are labeled as in A-H, but lower panels in each pair show the *unc-103e::mCherry* fluorescence used to confirm identity of muscle cells. Note that mCherry labeling of um1 is not visible in (J). (Q-V) Examples of GPCR::GFP expression in cells of the egg-laying system other than neurons and muscles. Images are labeled as in A-H, but lower panels in each pair show *ajm-1::mCherry* fluorescence. sp, ut, utse, uv3 and vt cells are identified by their outlines visualized with *ajm-1::mCherry*.

**Figure 5:**
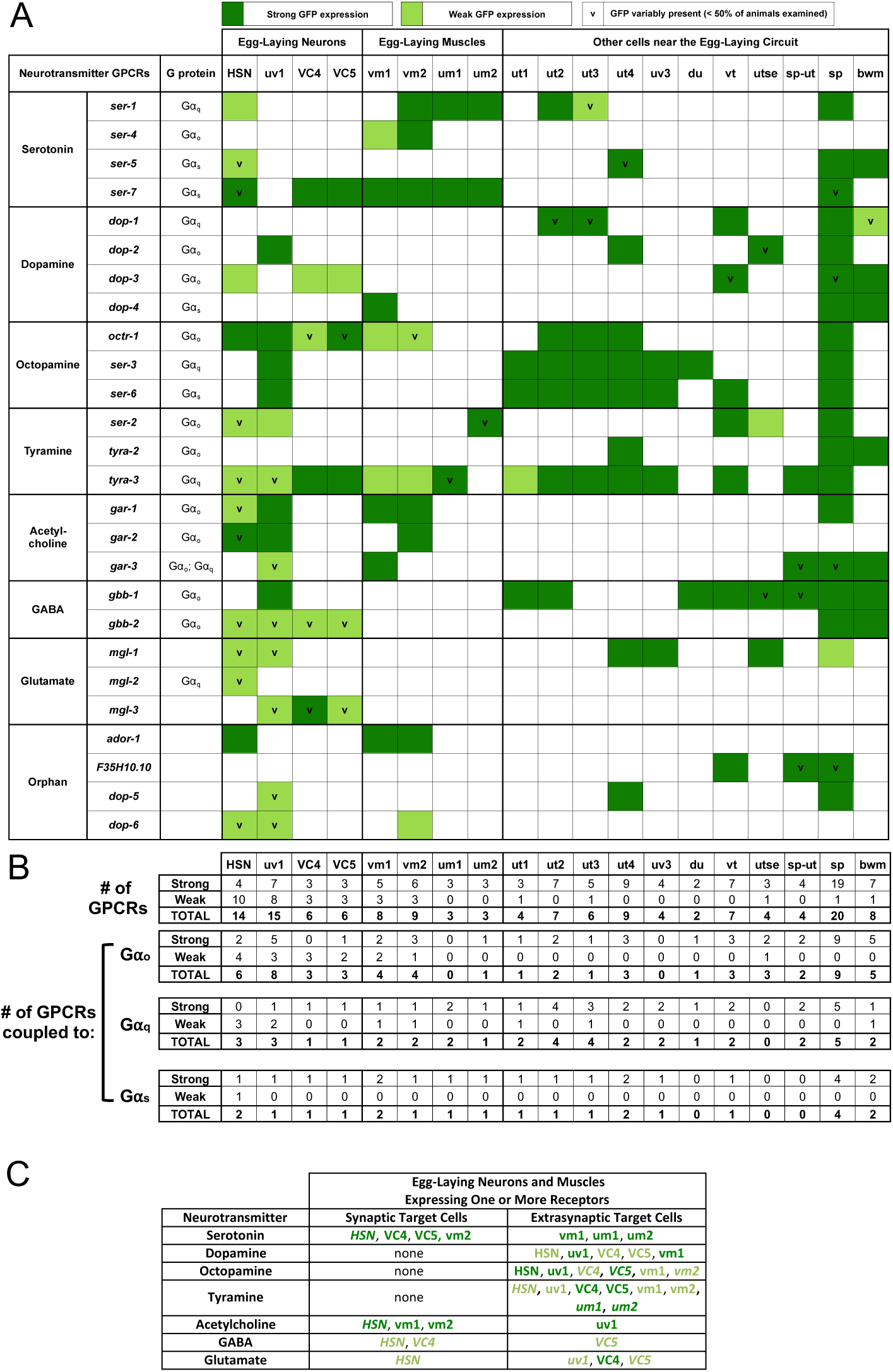
**Summary of Neurotransmitter GPCR::GFP Expression in the *C. elegans* Egg-Laying System** (A) Summary of neurotransmitter GPCR::GFP expression by cells in and near the egg-laying system. Dark green represents GFP expression that was on average strong (scored as +++, among brightest of any cell in the animal) or easily detectable (++). Light green represents weak GFP expression (+, just above background in at least two different animals). White indicates cells in which GFP was not reproducibly observed. “v” (for “variable”) denotes cells in which GFP detected in more than two but less than half of animals observed. Cell name abbreviations used here but not in prior figures - bwm: body wall muscles; sp-ut: spermathecal-uterine valve.. (B) Number of strongly and weakly expressed receptors in each cell type of the egg-laying system. Receptors with variable expression are included. Totals are shown broken down by the Gα protein thought to be activated by the receptors in each cell type when that information is known. (C) Neurons and muscles of the egg-laying system that express neurotransmitter GPCRs, indicating which cells have the corresponding neurotransmitter synaptically released onto them (synaptic target cells) or not (extrasynaptic target cells), based on the connectome (Albertson & Thomas, 1976; White et al., 1986; Xu et al., 2013) and neurotransmitter maps (Serrano-Saiz et al., 2013; Pereira et al., 2016; Gendrel et al., 2016). Cell names are dark/light green to indicate strong/weak receptor expression, while italics indicate receptor expression is variable.

Although 13 of the 26 neurotransmitter GPCRs were previously reported to be expressed in the egg-laying system, the vast majority of the expression we found was not previously reported (Figure S4A). In 20 instances we reproduced a published report of a receptor being expressed in a particular cell type, but in 119 additional instances we made a novel observation of a receptor being expressed in a cell type. In eight instances a published case of a receptor being expressed in a cell type was not reproduced in our data (Figure S4A).

While GFP reporters were often expressed strongly and reproducibly in a given cell type, we also to observed cases of weak and/or variable GFP expression (Figures 5 and S4B). This phenomenon is illustrated for *octr-1::gfp* expression in the HSN, VC4, and VC5 neurons in Figure S2. We collected data for all detectable expression so that future studies can investigate what types of expression of a GPCR::GFP reporter might reflect functional presence of the corresponding endogenous receptor.

We note that single-cell RNA sequencing (RNAseq) has the potential to map which cells express any gene, including the neurotransmitter GPCR genes we characterize here. Our data are far richer than the currently-available RNAseq data for the egg-laying system. Our work scores expression in 19 cell types in the egg-laying system (Figure 5) versus only four in the current RNAseq data (Taylor et al., 2020). As examples, we scored expression in vm1 and vm2 separately rather than aggregating all vulval muscle cell expression, and we included in our analysis cells other than neurons and muscles, which turn out to express many neurotransmitter GPCRs. Further, we measured variability in expression from animal to animal, and discovered weak expression below the detection threshold of current RNAseq data.

### Individual Cells Express Multiple Neurotransmitter GPCRs

Individual cells expressed up to 20 different neurotransmitter GPCRs (Figure 5). Neurons (examples in Figure 4A-H) each expressed 2-7 receptors at strong, reproducible levels and 4-10 additional receptors at weak and/or variable levels (Figure 5). Non-neuronal cells also expressed multiple neurotransmitter GPCRs strongly and reproducibly, but unlike neurons, these cells expressed only 0-3 additional receptors at weak and/or variable levels. Figure 4I-4P shows examples of GPCR::GFP expression in muscles cells, while Figure 4Q-4V shows examples of GPCR::GFP expression in non-neuronal, non-muscle cells of the egg-laying system.

The multiple receptors expressed on a single cell often couple to different G proteins (Figure 5B). Receptors that couple to Gα_q_ and Gα_s_ cooperate to activate neurons and muscles, while receptors that couple to Gα_o_ oppose such activation (reviewed in Koelle, 2018). Therefore, our results suggest individual cells of the egg-laying system have the ability to synthesize information received from multiple neurotransmitters and to compute an appropriate resulting level of activity by integrating signaling from multiple receptors and G proteins.

### Neurotransmitter GPCRs are Often Positioned to Receive Extrasynaptic Signals

As expected, neurotransmitter receptors were often expressed on cells that receive synapses from neurons releasing their activating neurotransmitter. More interestingly, receptors were also expressed on cells that do not receive any synapses at which the corresponding neurotransmitter is released (Figure 5C).

Serotonin receptors illustrate expression in cells positioned to receive synaptic versus extrasynaptic signals. In the egg-laying circuit, HSN neurons release serotonin to stimulate egg laying at synapses onto both the vm2 muscle cells and VC neurons (White et al., 1986; Collins and Koelle, 2013; Brewer et al., 2019). We observed that serotonin receptors are expressed, as expected, on these vm2 and VC postsynaptic cells (Figure 5). If we restrict our analysis only to strongly expressed receptors, vm2 cells express three serotonin receptors: the SER-1 and SER-7 receptors that stimulate egg laying by coupling to Gα_q_ and Gα_s_, respectively, and the SER-4 receptor that inhibits egg laying by coupling to Gα_o_ (Cho et al., 2000; Carnell et al., 2005; Dempsey et al., 2005; Carre-Pierrat et al., 2006; Hapiak et al., 2009; Gürel et al., 2012). The VC neurons express SER-7, which could mediate the ability of the HSN to excite the VCs (Collins et al., 2016). Finally, the HSN variably expresses SER-7, which might serve as an autoreceptor to allow the HSN to excite itself. The above instances of serotonin receptor expression all could mediate synaptic serotonin signaling.

Remarkably, our data show seven additional instances in which serotonin receptors are strongly and reproducibly expressed by cells of the egg-laying system that do not receive synapses from the HSN or any other serotonergic neurons (Figure 5). For example, the vm1 and um1/2 egg-laying muscle cells, which contract with the vm2 muscle cells to cause egg laying, all express serotonin receptors. Therefore, serotonin released by HSN may travel extrasynaptically to the vm1 and um1/2 cells to signal through serotonin receptors on these cells to help induce their activity and egg-laying events.

Acetylcholine, GABA, and glutamate, like serotonin, each have GPCRs expressed on cells onto which these neurotransmitters are synaptically released. In addition, acetylcholine, GABA and glutamate each have GPCRs expressed on neurons, muscles, and/or epithelial cells of the uterus that could receive these signals extrasynaptically (Figures 5A and 5C).

No cells of the egg-laying system receive synapses from neurons releasing dopamine, octopamine, or tyramine, yet receptors for each are expressed in the egg-laying system (Figure 5). For example, each of the four dopamine receptors DOP-1 through DOP-4 is strongly and reproducibly expressed in at least one cell type of the egg-laying circuit (Figure 5A). Indeed, endogenous dopamine has been shown to promote egg laying (Nagashima et al., 2016) and exogenous dopamine has been reported to both promote and inhibit egg-laying (Schafer and Kenyon, 1995; Vidal-Gadea et al., 2012). Similarly, application of octopamine and tyramine have been shown to inhibit egg laying (Horvitz et al., 1982; Alkema et al., 2005; Collins et al., 2016), and as in the case of dopamine, these effects may occur via extrasynaptic signaling through the receptors for these neurotransmitters we found expressed in the egg-laying system.

### Neurotransmitter GPCRs are Expressed by Cells other than Neurons and Muscles

In addition to being expressed in neurons and muscles (Figure 4A-4P), GPCR::GFP reporters were also frequently expressed on other cell types not previously known to respond to neurotransmitters (Figures 4Q-4V and 5A and 5B). Such cells include the uterine toroid epithelial cells ut1-ut4 that form a tube that holds unlaid eggs (Figures 2F, and 2G), the vulval toroid (vt) epithelial cells that form the opening through which eggs are laid (Figures 2F and 2G), and the spermatheca that holds sperm and through which eggs pass to be fertilized (Figures 2F and 2G).

The presence of neurotransmitter receptors on epithelial cells has precedent in mammals, as serotonin receptors are found on epithelial/endothelial cells gut (Berger et al., 2009; Hoffman et al., 2012) and blood vessels (Ullmer et al., 1995; Berger et al., 2009). The presence of receptors on the uterine toroid epithelial cells may indicate that these ut cells receive neurotransmitter signals to aid in pushing eggs out of the uterus (Newman, 1996). Neurotransmitter receptors on the spermatheca may regulate its stretch and contraction before ovulation (Wirshing and Cram, 2017), particularly since downstream effectors of the G protein Gα_q_ have been shown to be involved in ovulation (Clandinin et al., 1998).

### Knockout Mutants Suggest Neurotransmitter GPCRs Function Redundantly

We analyzed knockout mutants for each of the receptors for changes in the frequency of egg-laying behavior. One assay of this behavior counts unlaid eggs in a worm (Figure 6A). A wild-type worm both produces and lays eggs, accumulating ∼18 unlaid eggs in its uterus at steady state. Mutants with a hyperactive or inactive egg-laying circuit continue to produce eggs but accumulate fewer or more unlaid eggs, respectively (Chase and Koelle, 2004). A second assay of egg-laying behavior scores the developmental stage of just-laid eggs. Eggs in wild-type animals undergo multiple rounds of cell divisions while they wait in the uterus to be laid, but eggs of hyperactive egg-layers are laid shortly after being fertilized and often at earlier stages of development, with fewer than eight cells (Figure 6A).

**Figure 6:**
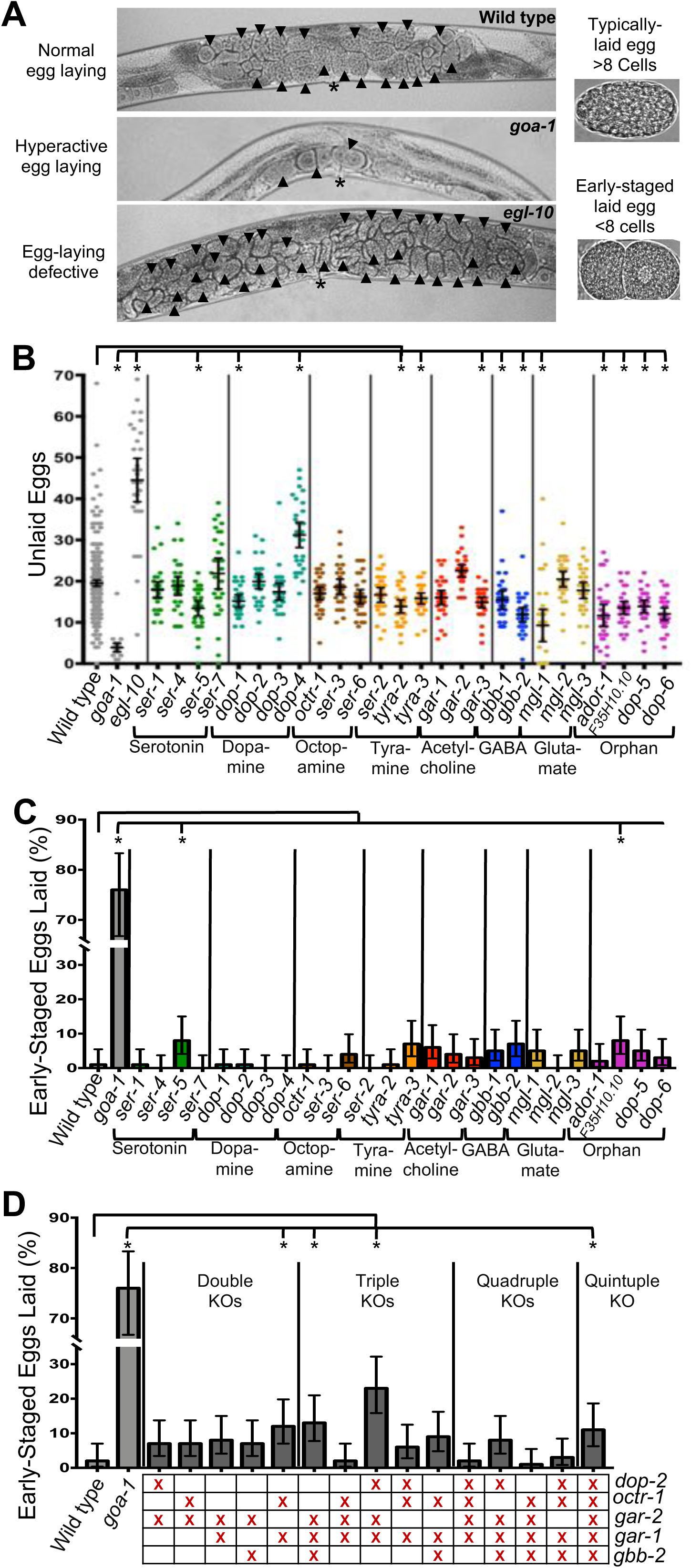
**Neurotransmitter GPCRs Knockout(s) Rarely Reveal Egg-Laying Defects** (A) Micrographs of animals with egg-laying phenotypes. Wild-type adults (top) typically accumulate 15-20 unlaid eggs, while hyperactive egg-laying mutants like *goa-1* (second from top) accumulate fewer unlaid eggs and egg-laying defective mutants like *egl-10* (third from top) accumulate more unlaid eggs. Eggs from wild-type animals have developed to the ∼100-cell stage by the time they are laid (top right), while eggs from hyperactive egg-laying mutants are laid earlier, often with fewer than eight cells (bottom right). (B) Number of unlaid eggs for neurotransmitter GPCR single knockout strains. *egl-10(md176)* and *goa-1(n1134)* served as controls (these control data are re-plotted in Figures 7C and S5B). n ≥ 30 for each strain. Asterisks indicate statistical significance (p<0.05) compared to the wild type using one-way ANOVA with Bonferroni correction for multiple comparisons. Error bars indicate 95% confidence intervals. Here and in other graphs in this work, different colors are used to plot data for receptors for different neurotransmitters, or for the orphan GPCR class. (C) Percent of early-stage eggs laid by neurotransmitter GPCR gene single knockout strains. Here and in (D), *goa-1(n1134)* is used as a control hyperactive egg-laying mutant (this control data is re-plotted in Figures 6D and S6A). n=100 eggs per strain. Asterisks indicate statistical significance (p<0.05) compared to the wild type using the Fisher’s exact test. The Wilson-Brown Method was used to determine the 95% C.I. for binomial data. (D) Percent of early-stage eggs laid by strains carrying combinations of knockouts in genes encoding G⍺_o_-coupled neurotransmitter GPCRs. *goa-1(n1134)* is used as a control hyperactive egg-laying mutant. Red “X” identifies the receptors knocked out in a given strain. Statistical analysis was as in (C).

Single knockouts of neurotransmitter GPCRs showed at most modest changes in the number of unlaid eggs accumulated, with the exception of the *dop-4* knockout (Figure 6B). The significance of the *dop-4* phenotype is dubious since we were unable to rescue it by transgenically re-expressing DOP-4 (Figure S5A). Twelve GPCR knockouts accumulated significantly fewer unlaid eggs than did the wild type (Figure 6B); however, only two of these laid significantly more early-stage eggs than did the wild-type control, suggesting that the other ten may have accumulated fewer unlaid eggs simply due to decreased egg production, not a change in egg laying. Even the two that laid significant early-stage eggs did so levels far below the 50% early-stage egg threshold considered biologically significant (Bany et al., 2003).

Because the *goa-1* mutant for the G protein Gα_o_ has a strong hyperactive egg-laying phenotype (Figure 6), we hypothesized that the effects of knockouts for receptors that signal through this G protein were masked by functional redundancy. Thus, we crossed together knockout mutations for G⍺_o_-coupled neurotransmitter GPCRs to see if this would reveal a strong hyperactive-egg-laying phenotype. We generated a series of combination knockouts, from double knockouts to a quintuple knockout, but none of these revealed a hyperactive egg-laying phenotype approaching the strength of the *goa-1* mutant (Figure 6D). One possibility is that Gα_o_ signals through even more Gα_o_-coupled receptors, such as neuropeptide receptors (Ringstad and Horvitz, 2008), which would need to be knocked out to phenocopy the Gα_o_ mutant.

### Overexpression of Neurotransmitter GPCRs Reveals Egg-Laying Defects

We next analyzed strains that overexpress individual neurotransmitter receptors, reasoning that this could result in gain-of-function phenotypes that might reveal receptor functions, even if individual receptors function redundantly with each other. Our high-copy GPCR::GFP transgenes should overexpress neurotransmitter receptors. The 20 C-terminal::GFP reporters overexpress a receptor with GFP fused to its C-terminus, while the four SL2::GFP reporters overexpress a receptor without any such modification (Figure S1). Two of our GPCR::GFP transgenes (for SER-4 and MGL-2) were constructed for technical reasons in a manner that would not cause receptor overexpression, so we did not analyze overexpression of these two receptors.

We piloted the neurotransmitter receptor overexpression strategy with the serotonin receptor SER-1, which had previously been reported to stimulate egg laying by acting partially redundantly with the serotonin receptor SER-7 (Hapiak et al., 2009). In the context of this experiment, we refer to the *ser-1::gfp* transgene as “*ser-1(oe)*” to indicate that the *ser-1* gene is <u>o</u>ver<u>e</u>xpressed. The experimental strategy is schematized in Figure 7A: we expected SER-1 overexpression to increase serotonin stimulation of egg laying and result in hyperactive egg laying, and that knocking out serotonin biosynthesis would suppress this effect since the overexpressed SER-1 would no longer be activated by serotonin. As expected, the *ser-1(oe)* strain displayed a strong hyperactive egg-laying phenotype (Figure 7B). The same phenotype was also observed in animals carrying an extrachromosomal *ser-1(oe)* transgene and in two independent chromosomal integrants for the *ser-1(oe)* transgene (Figure S6B), suggesting the phenotype was not an artifact of chromosomal integration. The hyperactive egg-laying phenotype of *ser-1(oe)* was completely suppressed in animals lacking TPH-1 (Figure 7B), an enzyme required for serotonin biosynthesis (Sze et al., 2000), suggesting that the effect of overexpressing SER-1 is due to increased serotonin signaling. These results were consistent with the previous data suggesting that SER-1 is one of two partially redundant receptors through which serotonin signals to stimulate egg laying (Hapiak et al. 2009).

**Figure 7:**
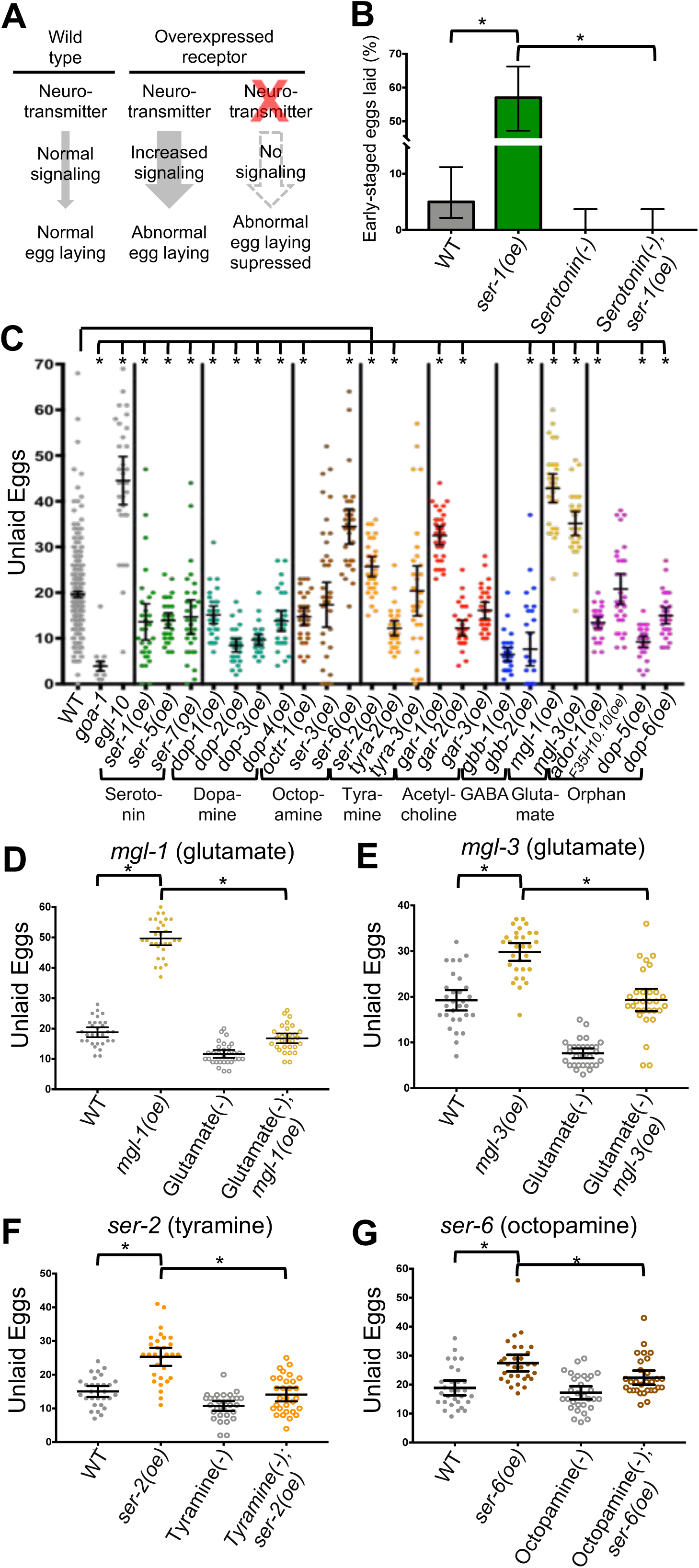
**Neurotransmitter GPCRs Overexpressors Reveal Strong Egg-Laying Defects** (A) Strategy to reveal and validate egg-laying defects caused by increased signaling through overexpressed neurotransmitter GPCRs. (B) Overexpression of the serotonin GPCR SER-1 (*ser-1(oe)*) causes hyperactive egg-laying that is suppressed by knocking out serotonin biosynthesis with a null mutant in the tryptophan hydroxylase enzyme TPH-1. n=100 eggs per strain. Error bars show 95% confidence intervals determined by the Wilson-Brown method. Asterisks indicate statistical significance (p<0.05) compared to the wild type using Fisher’s exact test. (C) Number of unlaid eggs for neurotransmitter GPCR gene overexpressor strains. *egl-10(md176)* and *goa-1(n1134)* served as controls. n ≥ 30 for each strain. Error bars, 95% confidence intervals. Asterisks indicate p<0.05 compared to the wild type using one-way ANOVA with Bonferroni correction for multiple comparisons. (D) Overexpression of the glutamate GPCR MGL-1 causes defective egg laying that is suppressed by knocking out glutamate signaling with a mutation in the vesicular glutamate transporter EAT-4. In (D-G), n ≥ 30 for each strain, error bars indicate 95% confidence intervals, and asterisks indicate p<0.05 using one-way ANOVA with a Tukey’s test to determine statistical significance for multiple comparisons. (E) Overexpression of the glutamate GPCR MGL-3 causes defective egg laying that is suppressed by knocking out glutamate signaling with a mutation in the vesicular glutamate transporter EAT-4. (F) Overexpression of the tyramine GPCR SER-2 causes defective egg-laying that is suppressed by knocking out the tyramine biosynthesis enzyme tyrosine decarboxylase (TDC-1). (G) Overexpression of the octopamine GPCR SER-6 causes defective egg laying that is suppressed by knocking out the octopamine biosynthesis enzyme tyramine beta-hydroxylase (TBH-1).

We examined all 24 neurotransmitter GPCR overexpressor transgenes and found six more instances of strong effects on egg laying. We could not validate two of these six cases: the GABA receptor GBB-1 and the octopamine receptor SER-3 each had one overexpressor transgene that resulted in changes to egg laying (Figure S6A and S6H), but these phenotypes were not reproduced with using other independent overexpressor transgenes (Figures S6C and S6H). However, the remaining four cases each suggest novel functional effects of neurotransmitter signaling in the egg-laying system, as described below.

Overexpression of the glutamate GPCRs *mgl-1(oe)* and *mgl-3(oe)* caused strong accumulation of unlaid eggs (Figure 7C). Extrachromosomal *mgl-1(oe)* and *mgl-3(oe)* transgenes also caused similar egg-laying defects (Figure S6F), and abolishing glutamate signaling by knocking out the vesicular glutamate transporter EAT-4 (Lee et al., 1999) completely suppressed the egg-laying defects of glutamate receptor overexpression (Figures 7D and 7E). In the egg-laying system, *mgl-1* is weakly and variably expressed in uv1 and HSN neurons but shows strong and consistent expression in specific epithelial cells of the uterus, while *mgl-3* shows its strongest (albeit variable) expression in the VC4 neuron (Figure 5). These results represent the first time glutamate signaling has been shown to affect *C. elegans* egg laying and provide a starting point for future analysis of this issue.

Overexpression of the octopamine receptor SER-6 caused accumulation of unlaid eggs (Figure 7C). We saw similar phenotypes using other extrachromosomal and chromosomally-integrated *ser-6(oe)* transgenes (Figure S6D), and could suppress this egg-laying defect by knocking out the octopamine biosynthetic enzyme TBH-1 (Alkema et al. 2005; Figure 7G). Previous studies had shown that exogenous octopamine inhibits egg-laying behavior (Horvitz et al., 1982) but the mechanism and receptor(s) mediating this effect had remained unknown. SER-6, along with other octopamine receptors, is strongly expressed in the uv1 neuroendocrine cells and in the epithelial cells of the uterus (Figure 5). To date, the uterine epithelial cells have not been implicated in regulation of egg laying. However, given that SER-6 is coupled to the excitatory G protein Gα_s_, we hypothesize that SER-6 could activate the uv1s, which in turn inhibit egg laying (Jose et al., 2007; Banerjee et al., 2017).

Overexpression of the acetylcholine receptor GAR-1 similarly caused accumulation of unlaid eggs (Figure 7C). We reproduced this defect with an extrachromosomal *gar-1(oe)* transgene (Figure S6G), but could not attempt to further validate this effect since a null allele for the acetylcholine biosynthetic enzyme is lethal (Rand, 1989). Inhibition of egg laying via GAR-1 signaling would be interesting as acetylcholine has been shown to both activate and inhibit egg laying (Bany et al. 2003; Schafer, 2006). GAR-1 is expressed principally in the uv1 neuroendocrine cells and the vm1/2 vulval muscles (Figure 5). As a Gα_o_-coupled receptor, GAR-1 is expected to reduce neural/muscle activity. Thus GAR-1 could activate egg laying by inhibiting the uv1 cells, which in turn inhibit egg laying, and GAR-1 could inhibit egg laying by inhibiting the vm1/2 muscles, which execute egg laying.

The final receptor that displayed a validated egg-laying defect when overexpressed was the tyramine receptor SER-2. The integrated *ser-2(oe)* transgene caused a mild retention of unlaid eggs (Figure 7C) that was reproduced with an extrachromosomal *ser-2(oe)* transgene (Figure S6I) and suppressed when we knocked out the tyramine biosynthetic enzyme TDC-1 (Alkema et al. 2005; Figure 7F). Tyramine was previously known to inhibit egg laying (Alkema et al., 2005; Collins et al. 2016). SER-2 is weakly and/or variably expressed in neurons (HSN, uv1) and muscle cells (um2) of the egg-laying system, but strongly expressed in certain non-neuronal/non-muscle cells, such as epithelial cells of the vulva (Figure 5). SER-2 is coupled to the inhibitory G protein Gα_o_, and we hypothesize that it could inhibit egg laying by acting in the HSN and um2 cells that act to stimulate egg laying.

### Rich and Functionally Robust Neurotransmitter GPCR Signaling Controls Neural Circuit Activity

This study provides for the first time analysis of the expression and functions of every neurotransmitter GPCR in a model neural circuit. The *C. elegans* egg-laying circuit was already one of the most intensively studied small neural circuits, yet this study provides new insights into this circuit that may apply to neural circuits in general.

The first insight is that there may be far more neurotransmitter signaling within neural circuits than had been suggested by previous work. All cells in the egg-laying system express multiple neurotransmitter receptors, and each cell type expresses its own unique combination of up to 20 such receptors. Cells of the egg-laying circuit express receptors for neurotransmitters not previously implicated in control of egg laying, such as glutamate and GABA. Cells previously thought to have merely structural roles, such as the epithelial cells that make up the uterus and vulva, also express neurotransmitter receptors. Expression and function of neurotransmitter receptors in cells other than neurons and muscles has not been widely studied, yet our results suggest this phenomenon may be widespread.

A second insight is that circuits are remarkable robust systems due to an apparent high level of redundancy and compensation among neurotransmitter GPCRs. While mutations that affect downstream signaling by the neural G proteins Gα_o_, Gα_q_, and Gα_s_ each show dramatic effects on egg laying (Schafer, 2006; Koelle, 2018), no single neurotransmitter GPCR receptor knockout causes significant defects. The receptor knockout strains we analyzed in our study are grossly healthy and move relatively normally (data not shown), suggesting that functional redundancy of neurotransmitter GPCRs extends beyond the egg-laying system. We found that overexpressing neurotransmitter GPCRs generates gain-of-function phenotypes that can reveal receptor functions despite the issue of redundancy. This strategy reveals novel potential functions of dopamine, octopamine, acetylcholine, GABA glutamate, and orphan receptors in controlling the egg-laying circuit. Receptor overexpression may similarly be useful in revealing redundant neurotransmitter signaling functions in other circuit.

Our results generate a rich set of hypotheses to direct future studies of the egg-laying circuit. For example, prior work has shown that egg laying is activated when the HSN releases a combination of serotonin and the neuropeptide NLP-3 (Brewer et al., 2019). These two signals act partially redundantly to turn on the egg-laying circuit, with serotonin itself acting through partially redundant receptors (Hapiak et al., 2009). The multiple layers of redundancy in this circuit activation step leads to circuit activity being highly robust, but they also complicate efforts to understand the mechanism of circuit activation. Our results show that the serotonin receptors that activate the circuit, principally SER-1 and SER-7 with SER-5 playing a lesser role (Hapiak et al., 2009), are each expressed in different overlapping sets of cells in the egg-laying system. SER-1 and SER-7 are both expressed in the vm2 muscle cells, whose serotonin activation is crucial for executing egg laying (Brewer et al. 2019). SER-7 is also expressed in the VC neurons, which synapse onto vm2. The HSNs activate the VCs, which fire every time an egg is laid (Collins et al., 2016), and our findings suggest the hypothesis that HSNs activate the VC neurons via the SER-7 serotonin receptor. We also found that the SER-1, SER-5, and SER-7 receptors are expressed on the HSNs themselves, suggesting the hypothesis that serotonin from the HSN could act in an autocrine feedback loop to maintain HSN activity. Prior work has shown that the uv1 cells release tyramine and a combination of neuropeptides (Collins et al., 2016; Banerjee et al., 2017), with these signals acting redundantly to inhibit the egg-laying circuit, and that tyramine itself acts through multiple redundant receptors. Our results show which cells of the circuit express each of the three tyramine receptors, setting the stage for studies to tease out the redundant mechanisms by which the circuit is inactivated.

The strategies and tools applied in this work for analyzing neurotransmitter signaling in the egg-laying circuit can be applied to any neural circuit in *C. elegans* or other organisms to generate sets of testable hypotheses for how signaling among the cells of a neural circuit control its activities and function.

## Acknowledgements

We thank Mihail Sarov and Susanne Hasse for providing a number of newly-recombineered GFP reporter fosmid clones, Eviatar Yemini and Oliver Hobert for the *mgl-2(7.9kb)::gfp* plasmid, and Kevin Collins and Helge Groβhans for the *ida-1::mCherry* and *ajm-1::mCherry* transgenes, respectively. Strains were provided by the *C. elegans* National BioResource Project of Japan and by the CGC, funded by the NIH Office of Research Infrastructure Programs (P40 OD010440). We thank the information resource WormBase. R.W.F. was supported by a Paul and Daisy Soros Fellowship and by NIGMS T32GM007223. K.W. was supported by the Yale STARS II program and E.Y.W. was supported by a Yale College Dean’s Research Fellowship. This work was supported by NIH grants NS036918 and NS086932 to M.R.K.

## Author Contributions

Conceptualization – R.W.F., K.W., M.R.K.; Methodology – R.W.F., K.W., D.M., M.R.K.; Formal Analysis - RW.F., K.W.; Investigation – R.W.F., K.W., E.Y.W., N.C.; Resources – R.W.F., K.W., E.Y.W, D.M., A.O., J.P., S.K.; Data Curation – R.W.F.; Writing – Original Draft. – R.W.F.; Writing - Review and Editing – R.W.F., M.R.K.; Visualization – R.W.F., K.W., E.Y.W., N.C.; Supervision – R.W.F., M.R.K.; Project Administration – M.R.K.; Funding Acquisition – M.R.K.

## STAR Methods

KEY RESOURCES TABLE

### CONTACT FOR REAGANT AND RESOURCE SHARING

Some *C. elegans* strains generated in this work will be made available via the *Caenorhabditis* Genetics Center (CGC). Further information and requests for other reagents and resources should be directed to and will be fulfilled by the lead contact, Michael R. Koelle.

### EXPERIMENTAL MODEL AND SUBJECT DETAILS

#### Strains and culture

*C. elegans* strains used in this study were maintained at 20°C on standard nematode growth media (NGM) seeded with OP50 strain of *Escherichia coli* as their food source. Null alleles for neurotransmitter receptors and biosynthetic enzymes were obtained through *Caenorhabditis* Genetics Center (CGC) the Japanese National BioResource Project (NBRP), or as otherwise indicated. Null alleles were backcrossed 2-10x to N2 (wild type), as indicated. New strains were constructed using standard procedures and genotypes were confirmed by PCR or sequencing. Extrachromosomal array transgenic strains for neurotransmitter GPCRs were generated through microinjection and at least one independent line was kept. Extrachromosomal strains for neurotransmitter GPCRs were chromosomally integrated through UV/TMP mutagenesis and at least one integrant was kept. Integrated strains were backcrossed 2-4x to N2 (wild type) as indicated. Young adult hermaphrodite animals were used for confocal microscopy. For egg-laying assays, animals were staged 40 hours post-L4. A complete list of all of the *C. elegans* strains used in this paper are provided in the Key Resources Table, with additional details found in Table S2.

### METHOD DETAILS

#### Molecular Biology

##### Isolating and sequencing recombineered fosmids

Recombineered transgenes were received from TransgeneOme (transgeneome.mpi-cbg.de) in EPI300. Clones were recovered and DNA was isolated as described by Sarov et al., (2012) with some modifications. Non-clonal cell populations carrying the constructs were streaked onto triple antibiotic plates (15µg/mL chloramphenicol, 100µg/mL streptomycin, and 50µg/mL cloNAT) and individual colonies were inoculated for 14-16hrs in LB broth with 1X Fosmid Autoinduction solution and the antibiotics. A subset of the transgenes (see below) do not contain the cloNAT resistance marker and therefore only chloramphenicol and streptomycin were used for their growth. Recombineered fosmid DNA was isolated using Qiagen Plasmid Mini Kits. The purified DNA was run on 0.4% agarose gels to determine if the recombineered fosmid was at the correct size. The GFP coding region of each fosmid was also sequenced to determine if the GFP tag was inserted at the correct position.

##### Microinjections

Recombineered transgenes were injected at 20-100ng/µL into *unc-119(ed3)* animals, or into *lin-15(n765ts)* animals along with a *lin-15* rescuing plasmid (pL15EK) at 50ng/µL, plus 25 ng/µl of *E. coli* DH5α genomic DNA that had been digested to an average length of ∼6 kb. Strains carrying recombineered transgenes are listed in the Key Resources table, with additional details found in Table S2.

##### Chromosomal Integrations

Extrachromosomal transgenic strains were chromosomally integrated following UV/TMP mutagenesis. At least 300 F1 brightly GFP positive animals were picked to individual plates. Thirty or more of the best of these F1 plates were selected based on the presence of F2 progeny that were > 75% non-multivulva (i.e. positive for the *lin-15* marker on the transgene) and brightly GFP positive. For each selected F1 plate, six F2 animals were placed on single plates. F2 homozygous integrants were identified as giving rise to true-breeding 100% non-Muv and 100% GFP positive strains. This procedure resulted in 1-7 independent integrants per transgene.

##### Transgenic reporter mCherry strains

All mCherry marker strains used in our study are chromosomally integrated. The pBR1 plasmid for *ida-1::mCherry* (a gift from Kevin Collins) was injected at 20 ng/µl with 50 ng/µl pL15EK plus 25ng/µl of *E. coli* DH5α genomic DNA digested to an average length of ∼6 kb into *lin-15(n765ts)* animals to generate extrachromosomal lines and chromosomally integrated. The *ida-1::mCherry* transgenic strain was used for cell identifications of the HSN, uv1s, VC4, and VC5 of the *C. elegans* egg-laying circuit. Independent integrants for *unc-103e::mCherry* previously generated in our lab (Collins and Koelle, 2013) were used for identification of the vulval (vm1 and vm2) and uterine (um1 and um2) muscles. The *ajm-1::mCherry* integrated strain was received from the Gottschalk lab. It labels the apical borders of epithelial cells and was used to identify the uterine toroid (ut1-4) cells, uv3 uterine ventral cells, vulval toroid (vt) cells, spermatheca-uterine (sp-ut) valve, spermatheca, dorsal uterine cell (du), and uterine seam (utse).

#### GPCR::GFP Transgenic Strains

##### C-terminal GFP constructs

Nineteen C-terminal GFP constructs listed in Table S1 were recombineered fosmid clones with GFP coding sequences inserted in the last coding exon of a receptor gene, between the codon for the C-terminal amino acid and the stop codon, as described in Sarov et al., (2012). Thirteen of these were constructed by Sarov et al. (2012), and details are available at the TransgeneOme web site (transgeneome.mpi-cbg.de). Five additional constructs (SER-1, SER-7, GAR-3, DOP-5, and DOP-6) were recombineered in an analogous fashion, except that for these constructs the *unc-119* and cloNAT resistant markers were not inserted into the final clone. The GFP reporter for *gbb-2* was recombineered in a similar fashion, except that GFP coding sequences were inserted in the middle of exon 14, such that GFP is inserted C-terminal to the amino acid Arg716. The most 3’ coding exons of the *gbb-2* gene have not been confidently determined, so this GFP insertion site was chosen as being 3’ to the coding sequences for the last transmembrane domain yet still confidently within the gbb-2 coding region.

##### SL2::NLS::GFP constructs

Microinjection of four C-terminal GFP transgenes (*dop-1::gfp*, *dop-4::gfp*, *gar-2::gfp*, and *F35H10.10::GFP*) resulted in GFP expression that was either weak or strong but localized to neural processes. In either case, cell bodies were not labeled with GFP adequately to allow identification of GFP positive cells. Using fosmid genomic clones (Table 1), these four receptor genes were recombineered to insert a an SL2 trans-splicing signal followed by GFP coding sequences directly after the GPCR gene stop codon, following the protocol of Tursun et al. (2009). The resulting transgenes express a single primary transcript that is processed into two separate mRNAs, one expressing the GPCR and the other expressing soluble GFP, allowing us to visualize GFP in the cell bodies.

##### Other constructs

Two GPCR::GFP transgenes could not be constructed by recombineering fosmids for technical reasons. The *ser-4::gfp* construct (Tsalik et al., 2003) is an integrated transgene that contains 4.1kb of *ser-4* promoter region 5’ of the start codon and fuses GFP coding sequences at an internal exon 3.7 kb downstream of the start codon. The *mgl-2::gfp* plasmid was a gift from Oliver Hobert (Yemini et al., 2019). It contains 7.9kb of the *mgl-2* promoter region 5’ upstream of the start codon followed by GFP coding sequences, which was injected to generate extrachromosomal lines and chromosomally integrated.

##### Isolation of GCPR::GFP restriction fragments from recombineered fosmid clones

Microinjection of recombineered fosmid clones for five GPCR genes (*dop-3::gfp*; *tyra-2::gfp*; *tyra-3::gfp*; *mgl-3::gfp*; *dop-5::gfp*) resulted in lethality or an absence of strong GFP expression in the extrachromosomal transgenic lines. To overcome these issues, 1-3µg of each recombineered transgene was digested with restriction enzymes to cut away other genes flanking the *gpcr::gfp* gene on the fosmid as described in Table 1. The digests were run on 0.4% agarose gels for >10 hours to purify the large fragments carrying the *gpcr::gfp* genes and gel extracted using the QIAEX II Gel Extraction Kit (Qiagen). The resulting product yielded a purified sample of 30ng/uL, a portion of which was run on an analytical 0.4% agarose gel to determine if product was at the expected size. To further confirm the correct fragment was purified, PCR primers were used to amplify the 5’ and 3’ flanking ends of the fragment and the products were analyzed on agarose gels. The purified restriction fragment transgenes were used for microinjections, resulting in satisfactory viability and GFP expression.

#### Confocal Imaging

Animals were on mounted on 2% agarose pads containing 120mM Optiprep (Sigma-Aldrich) to reduce refractive index mismatch (Boothe et al., 2017) on premium microscope slides superfrost (Fisher Scientific) and a 22×22-1 microscope cover glass (Fisher Scientific) was placed on top of the agarose pad. Animals were anesthetized using a drop of 150mM sodium azide (Sigma-Aldrich) with 120mM Optiprep. Z-stack confocal images of ∼100-130 slices (each ∼0.60*μ*m thick) images were taken on a Zeiss LSM 710 using a 40X objective lens. For at least 10 individual animals for each double-labeled transgenic strain, images of the mid-body where the egg-laying circuit lies were taken. Images were visualized using Vaa3D (Peng et al., 2010).

#### Behavioral Assays

For all egg-laying assays, unlaid eggs and early-staged eggs laid were quantified at 40 hours past the late L4 stage (Waggoner et al., 1998; Chase and Koelle, 2004; Collins and Koelle, 2013; Brewer et al., 2019).

### QUANTIFICTION AND STATISTICAL ANALYSES

Data graphs were made using Prism 7.0 software (GraphPad, La Jolla, CA). Scatter plots show individual data points (dots), mean (horizontal line), and error bars indicate 95% confidence intervals (C.I.) for the mean. N values are indicated in the figure legends and sample size for behavioral assays followed previous studies (Waggoner et al., 1998; Chase and Koelle, 2004; Collins and Koelle, 2013; Brewer et al., 2019).

Statistical significance was tested using one-way ANOVA with Bonferroni correction for multiple comparisons for the unlaid egg assay and Fisher’s exact test for the early-staged egg-laying assay. For the unlaid egg assay for the *dop-4* rescue experiment, validation of the neurotransmitter GPCR overexpressors, and the biosynthetic enzyme knockouts, the Tukey’s test was used to determine statistical significance. For the early-staged egg-laying assay, the Wilson-Brown Method was used to determine the 95% C.I. for binomial data. p ≥0.05 was considered not significant (ns) and p <0.05 (*) was considered significant.

### DATA AND SOFTWARE AVAILABILITY

All data, including confocal images, are available on request as electronic files.

---

**Figure S1:**
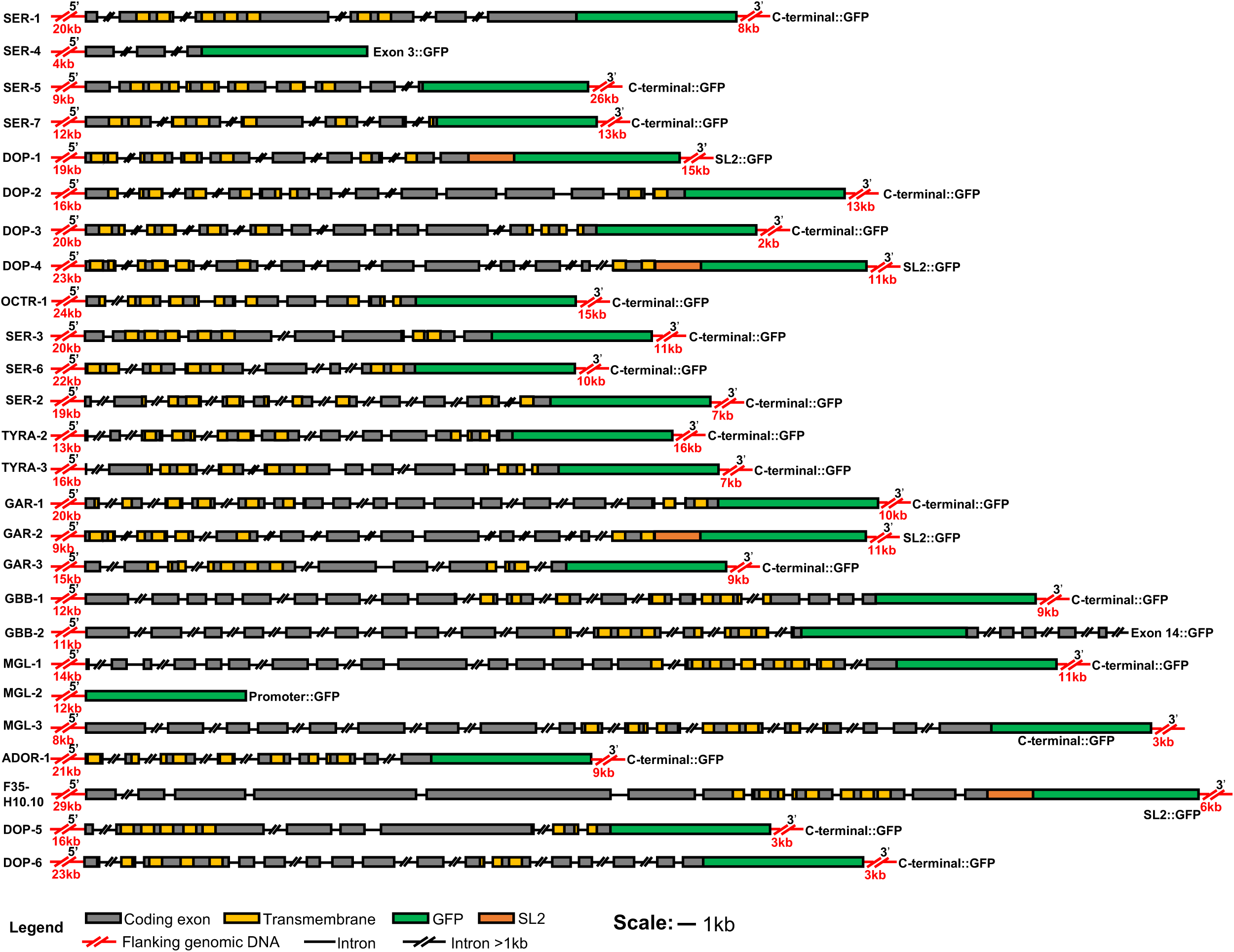
**GFP Reporters for 26 Neurotransmitter GPCRs, Related to Figure 1** Schematic of GFP reporters used in this study. For all reporters, the lengths of the 5’ promoter and 3’ downstream regions are indicated in kilobases (kb). Coding exons are indicated by gray boxes, predicted transmembrane domain coding regions by yellow boxes, GFP coding regions by a green box, and the SL2 trans-splicing signal region by an orange box. Flanking genomic DNA is indicated in red. Introns 1kb or less are drawn to scale with a black line. Introns greater than 1 kb are not drawn to scale and indicated with slashed black lines. C-terminal::GFP constructs were used for 19 neurotransmitter receptors and express a receptor with GFP fused to its C-terminus. SL2::GFP constructs were used for four neurotransmitter receptors: an SL2 trans-splicing signal followed by GFP coding sequences were inserted immediately downstream of the neurotransmitter GPCR stop codon so that such reporters co-expresses a neurotransmitter GPCR and GFP as separate proteins. Three transgenes were constructed in other manners. The *ser-4::gfp* transgene (Tsalik et al., 2003; Gürel et al., 2012) has GFP coding sequences fused to the third coding exon of *ser-4.* Repetitive sequences in the *ser-4* genomic DNA 3’ of this position prevented us from including the downstream regions of *ser-4* in the transgenes. The *gbb-2::gfp* reporter had GFP coding sequences inserted between two arginine codons in exon 14 of the *gbb-2* gene. The details of splicing downstream of exon 14 remain uncertain, preventing us from inserting GFP coding sequences more 3’ to this exon, The *mgl-2::gfp* reporter is a transcriptional fusion with a promoter fragment extending 7.9 kb 5’ of the *mgl-2* start codon inserted upstream of GFP coding sequences in the plasmid pPD955_75 (Addgene).

**Figure S2:**
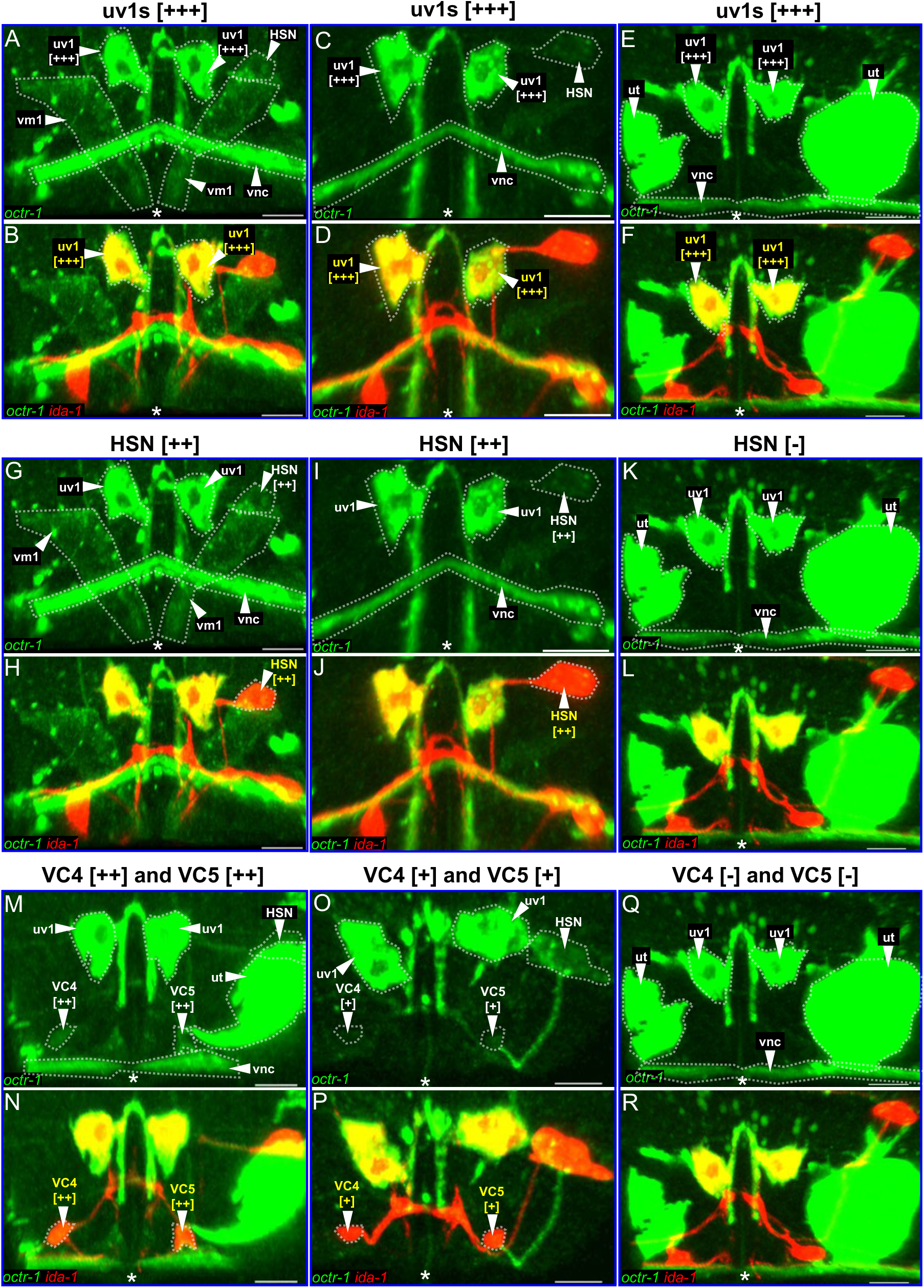
**Animal-to-Animal Variations in GFP Expression from a Chromosomally-Integrated *octr-1::gfp* Transgene, Related to Figure 3** (A-F) Strong expression of *octr-1::gfp* in uv1 cells is consistent from animal to animal. Confocal images of *octr-1::gfp; ida-1::mCherry* in the GFP channel (A, C, E) and GFP plus mCherry channel (B, D, F) for three different young adult animals. In (A, C, E) cells other than uv1 that express GFP are indicated by white text. GFP intensity in the cells of interest is indicated by +++ (among the strongest cells labeled), ++ (easily detectable), + (slightly above background). Labeling in (G-R) is analogous to that in (A-F). (G-L) Expression of *octr-1::gfp* in the HSN neurons varies from easily detectable to undetectable in different animals. The same confocal images as in (A-F) are shown but in these panels the labels for HSN include the GFP levels scored for this cell. GFP is easily detectable in two animals shown (G-I) but not detectable above background in a third (K and L). (M-R) Expression of *octr-1::gfp* in the VC neurons varies from moderate to undetectable in different animals. Labels for VC neurons indicate the GFP levels scored for these cells. Q and R are the same images shown in E, F, K and L but with new labels, while M-O are images not shown in previous panels. VC4/5 neuron GFP varies between animals, from easily detectable (M and N), to slightly above background (O and P), to undetectable (Q and R). In all panels, asterisks indicate position of the vulva; HSN: hermaphrodite-specific neuron; uv: uterine ventral cell; VC: ventral cord type C neuron; vm: vulval muscle; vnc: ventral nerve cord; scale bars: 10 microns.

**Figure S3:**
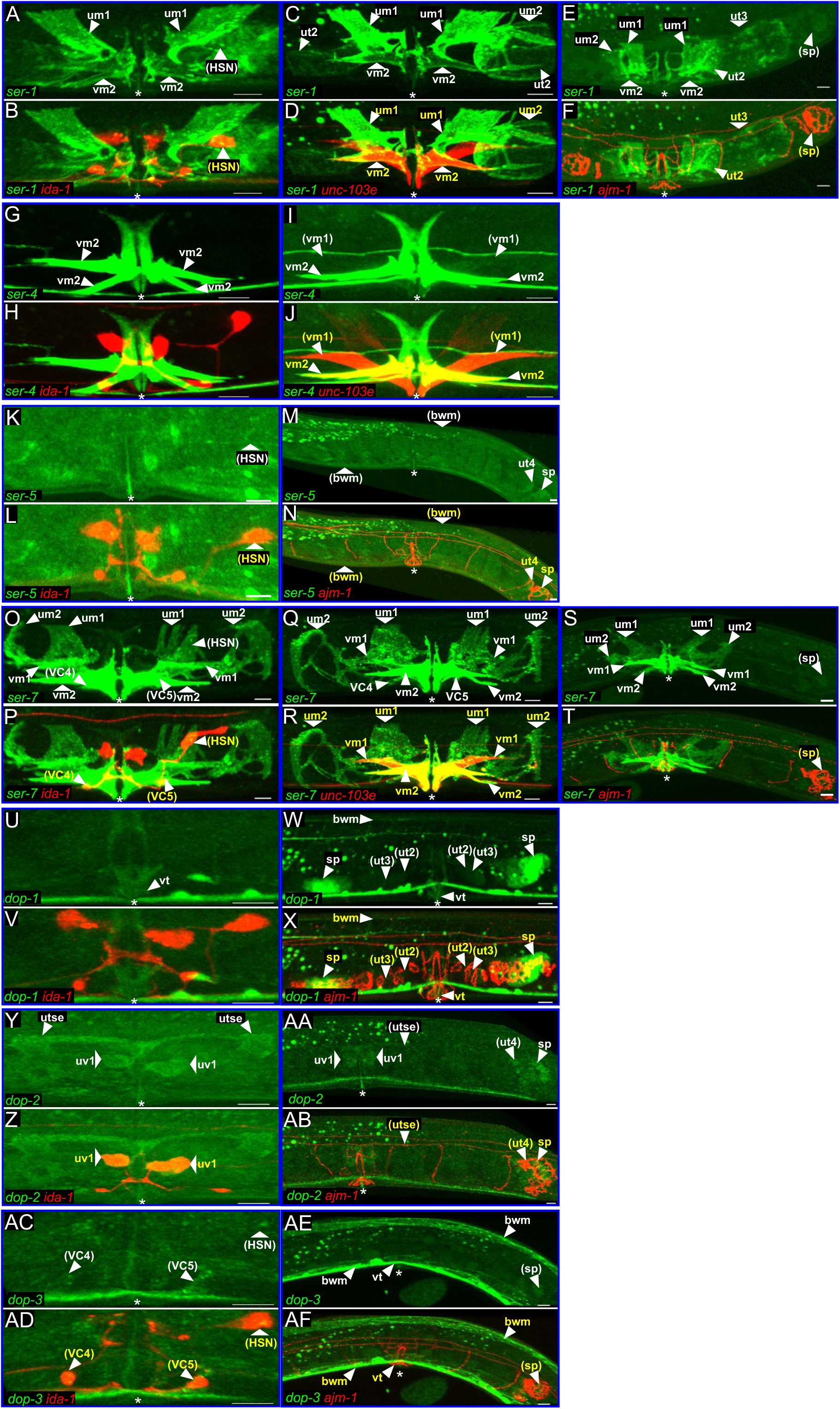
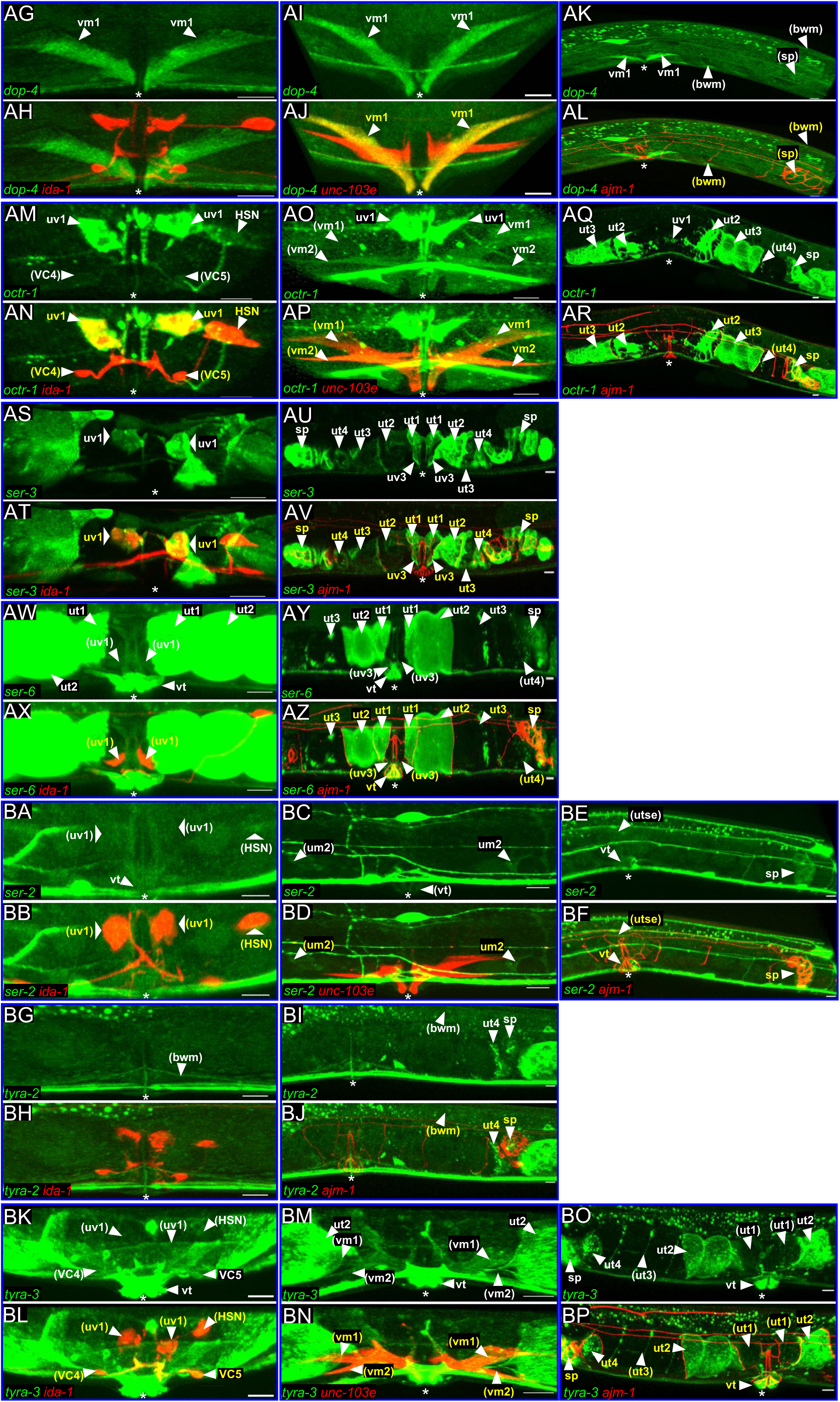
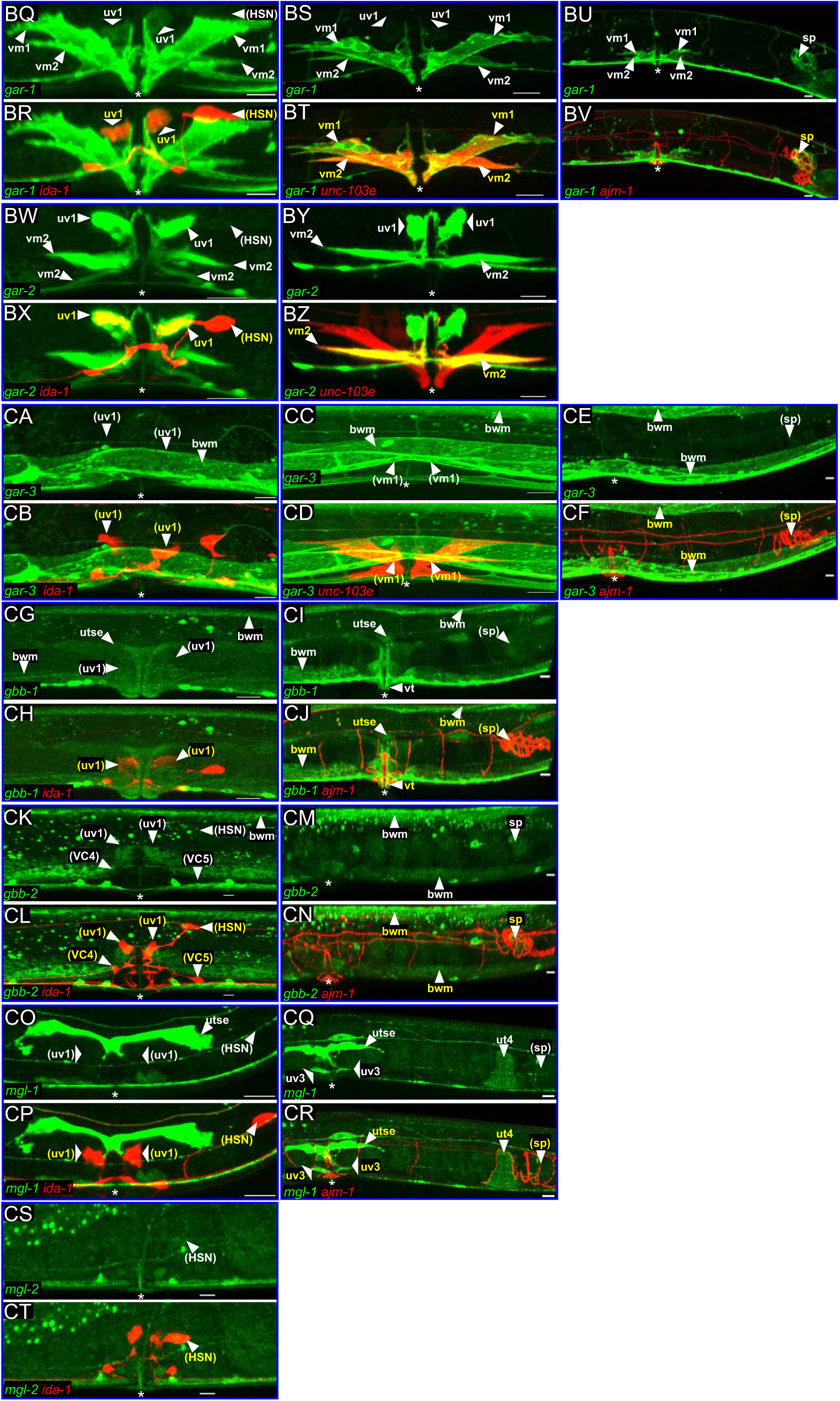
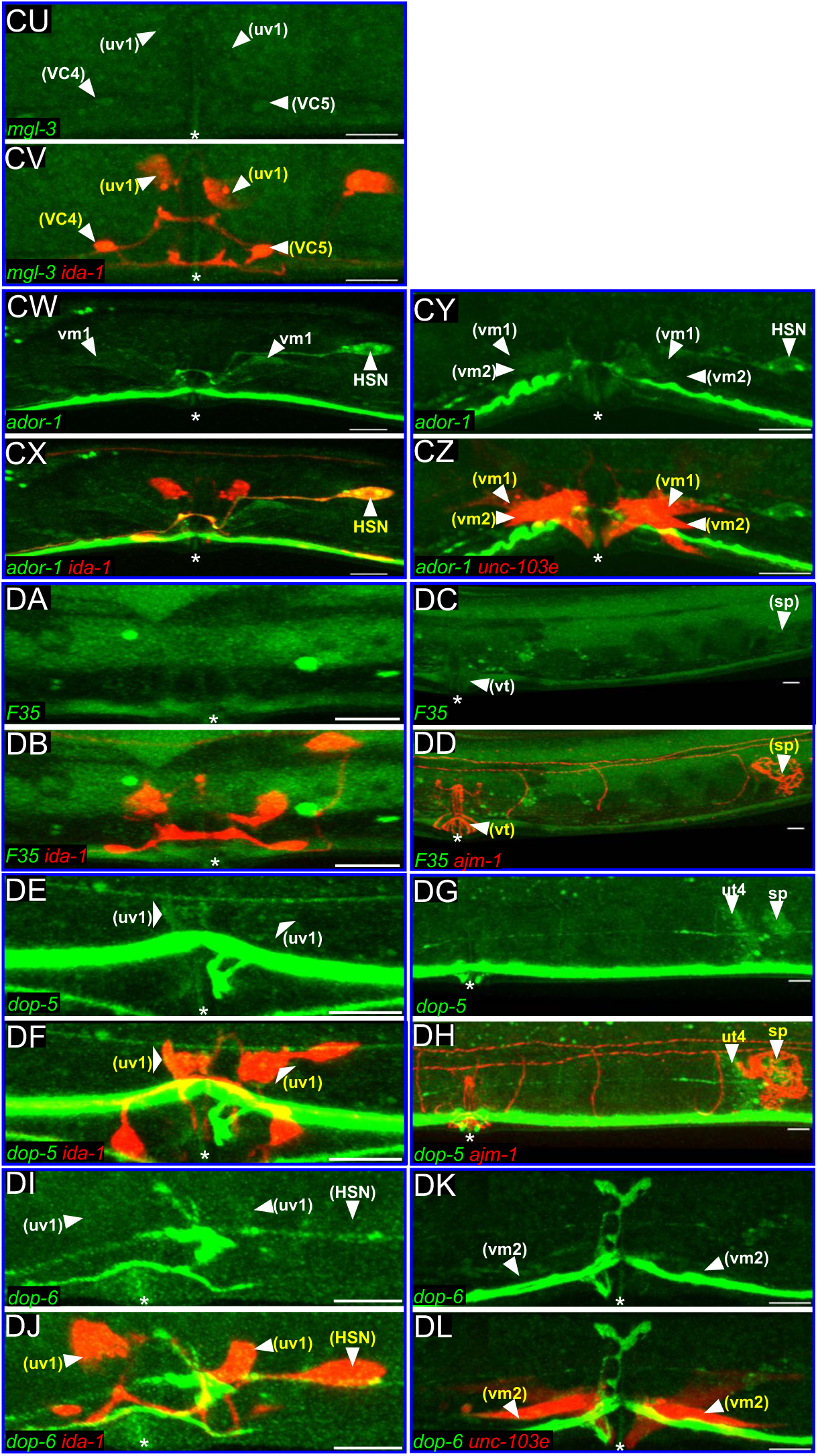
**Examples of GPCR::GFP Expression in the Egg-Laying System for All 26 Neurotransmitter GPCRs, Related to Figure 4** A larger subset of images than shown in Figure 4, selected from the 809 confocal images analyzed in this work. Actual analysis of GFP expression was carried out in three dimensions as shown in Video S1, and these additional two-dimensional images are shown for illustrative purposes. All images are presented in pairs, with the upper panel showing GFP fluorescence expressed from a reporter for the neurotransmitter GPCR named in green and the lower panel adding mCherry fluorescence for the marker transgene named in red. Upper panels use arrowheads and white cell names to indicate every cell type detectably expressing the GPCR::GFP reporter in our full data set. Lower panels use arrowheads and yellow cell names to indicate GFP-expressing cells that are also labeled by the mCherry marker. Cell names in parentheses indicate cells where GFP expression is not visible in the image as displayed, but can be detected in two or more such images from our full data set. White asterisks indicate the position of the vulva and all scale bars are 10 µm. Cell types are denoted by the following abbreviation: bwm: body wall muscles; du: dorsal uterine cell; HSN: hermaphrodite-specific neuron; sp: spermatheca; sp-ut: spermathecal-uterine valve; um: uterine muscle; ut: uterine toroid cell; utse: uterine seam cell; uv: uterine ventral cell; VC: ventral cord type C neuron; vm: vulval muscle; vt: vulval toroid cell.

**Figure S4:**
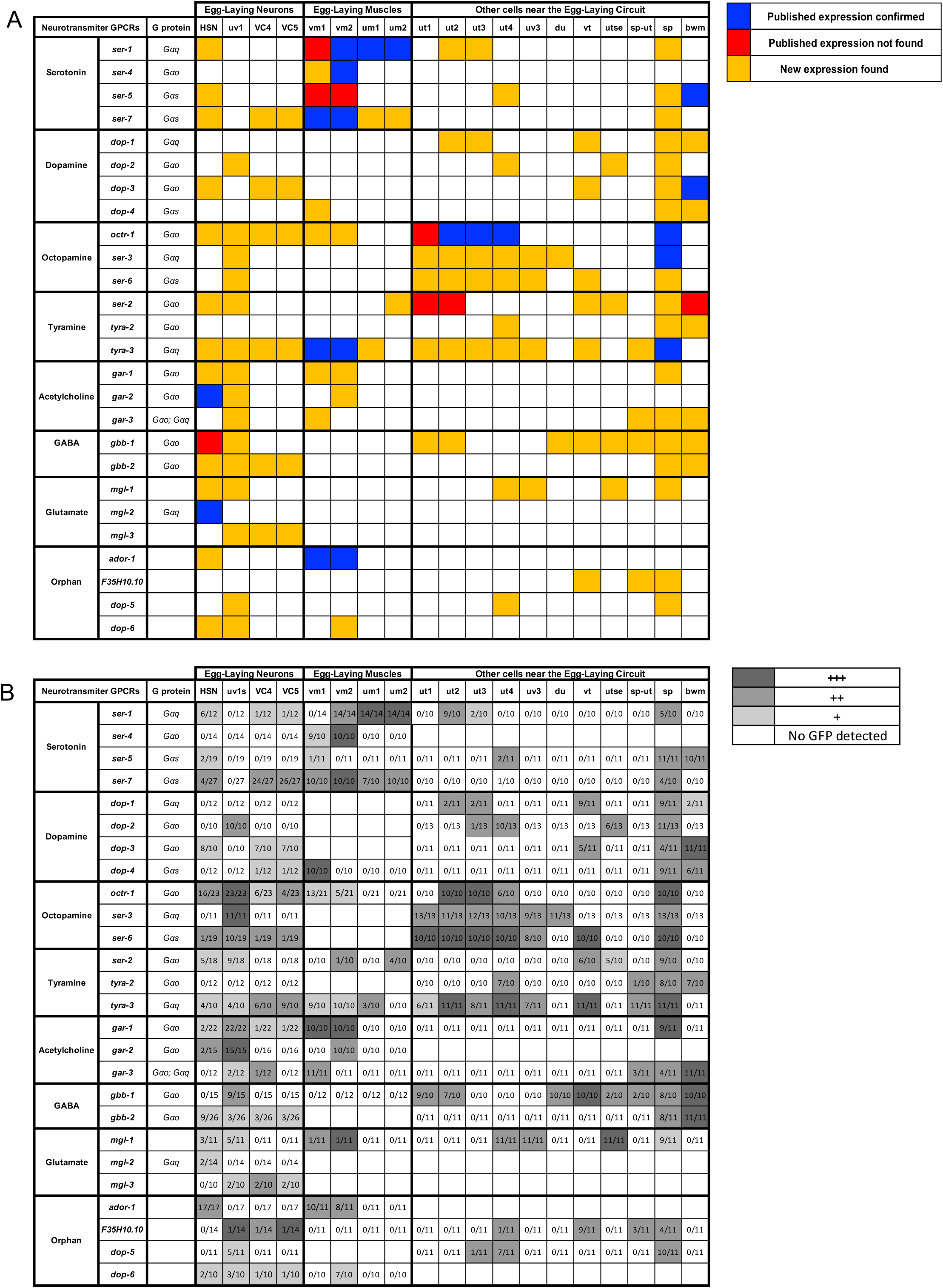
**Neurotransmitter GPCR Expressed Identified in this Work Compared to that Found in Previous Publications, Shown with Details of Variability, Related to Figure 5** (A) GPCR expression results from our study compared to previous findings. Blue indicates that we observed GPCR::GFP expression in a cell where expression of that receptor had previously been reporter, red indicates that we did not observe GPCR::GFP expression in a cell where such expression was previously reported, and orange indicates that we identified expression in a cell where no such expression had been previously reported. The previous studies that reported expression for 13 of the 26 neurotransmitter GPCRs in the egg-laying system were as follows: *ser-1*: Cho et al., 2000; Carnell et al., 2005; Carre-Pierrat et al., 2006; *ser-4*: Gürel et al., 2012; *ser-5*: Carre-Pierrat et al., 2006; Hapiak et al., 2009; *ser-7*: Carre-Pierrat et al., 2006; *dop-3*: Chase et al., 2004; *octr-1*: Wragg et al., 2007; *ser-3*: Suo et al, 2006; *ser-2*: Tsalik et al., 2003; Rex et al., 2004*; tyra-3*: Wragg et al., 2007; Bendesky et al., 2011; *gar-2*: Lee et al., 2000 ; *gbb-1*: Yemini et al., 2019 ; *mgl-2*: Yemini et al., 2019; and *ador-1*: Plummer, 2011. (B) Number of observations and strength of GFP expression in each cell type of the egg-laying system for each of the 26 GPCR::GFP reporters. Dark grey indicates cells in which GFP, when detected, was on average at the brightest levels (+++) for a reporter. Medium grey indicates cells in which GFP, when detected, was on average easily detectable (++). Light grey indicates cells that in which GFP, when detected, was on average only slightly above background (+). White indicates cells in which GFP was not observed in at least two animals. Numbers within a box in the form x/y indicate that GFP was detected x times in y animals examined for that cell type. Cell types are denoted by the following abbreviation: bwm: body wall muscles; du: dorsal uterine cell ; HSN: hermaphrodite-specific neuron; sp: spermatheca; sp-ut: spermathecal-uterine valve; um: uterine muscle; ut: uterine toroid cell; uv: uterine ventral cell; VC: ventral cord type C neuron; vm: vulval muscle; vt: vulval toroid cell.

**Figure S5:**
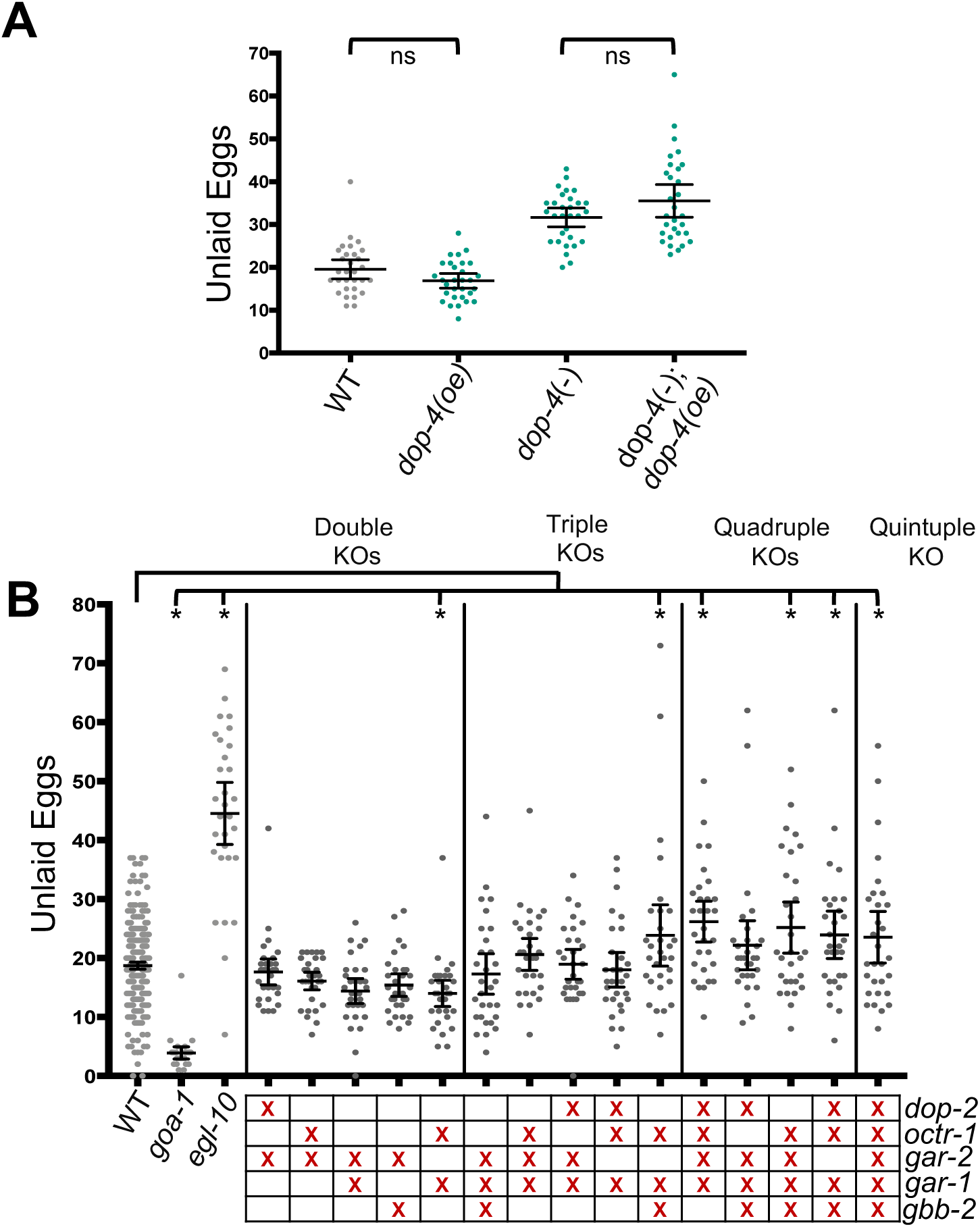
**Neurotransmitter GPCRs Knockout(s) Reveal No Significant Egg-Laying Defects, Related to Figure 6** (A) The increased number of unlaid eggs in the *dop-4* knockout strain cannot be rescued by introduction of a wild-type copy of the *dop-4* gene in the form of the chromosomally integrated *dop-4::SL2::NLS::gfp* construct. Statistical significance was tested using one-way ANOVA with a Tukey’s test to determine statistical significance for multiple comparisons for the unlaid egg assay. n ≥ 30 for each strain. p ≥0.05 was considered not significant (ns). (B) Combination knockouts of G⍺_o_-coupled neurotransmitter GPCRs expressed in the egg-laying system reveal no strong egg-laying defects in the accumulation of unlaid eggs in the uterus. In the table below graph, the red “X” signifies the combination of neurotransmitter G⍺_o_-coupled GPCR genes knocked out. *egl-10(md176) V* and *goa-1(n1134) I* served as controls for strong egg-laying defects. Statistical significance was tested using one-way ANOVA with Bonferroni correction for multiple comparisons for the unlaid egg assay. n ≥ 30 for each strain. p <0.05 (*) was considered significant.

**Figure S6:**
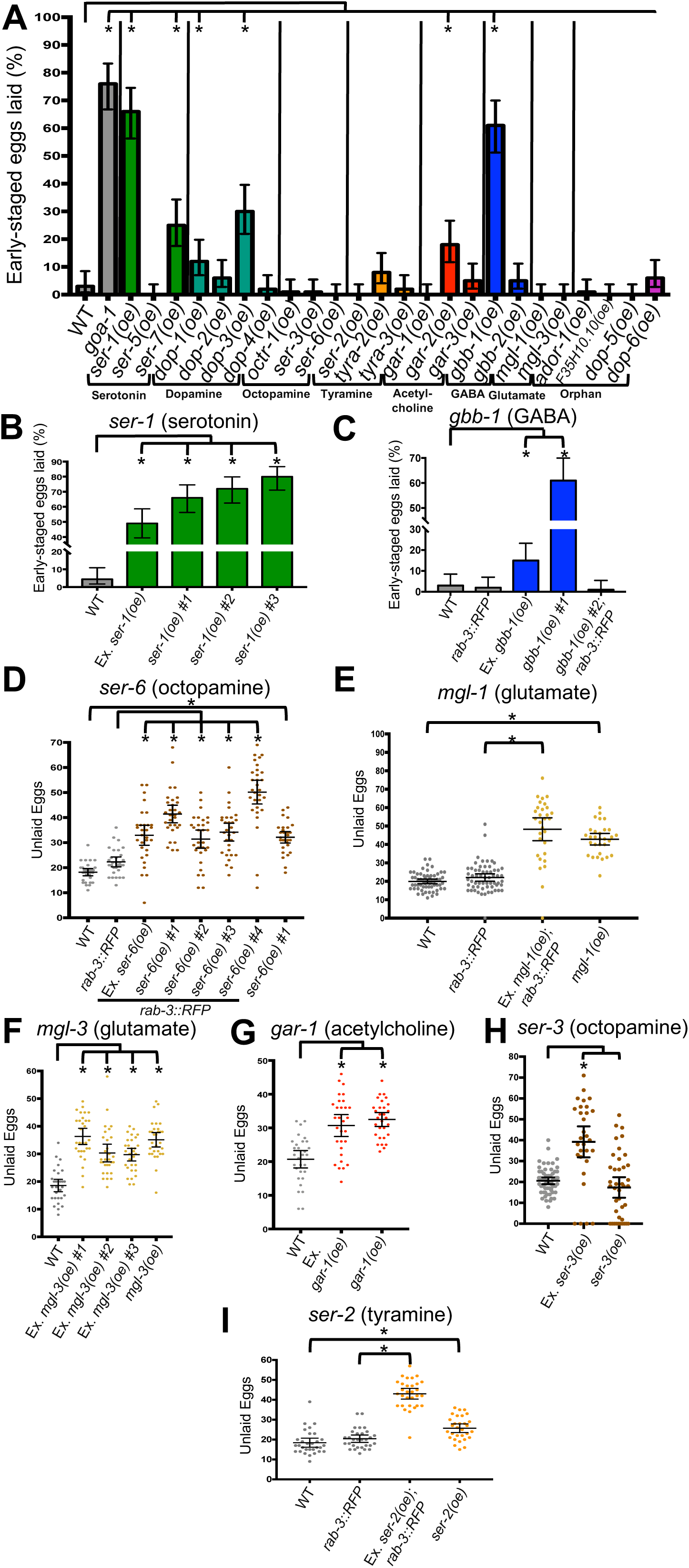
Egg-laying Defects in Neurotransmitter GPCRs Overexpressors Are Caused by Increased Neurotransmitter Signaling, Related to Figure 7 (A) Overexpression of two neurotransmitter GPCR genes, *ser-1* and *gbb-1*, results in a strong hyperactive egg-laying phenotype as seen by the high percentage of early-staged eggs laid. Bars are color coded for the neurotransmitter corresponding to teach receptor: dark green, serotonin; light green, dopamine; brown, octopamine; orange, tyramine; red, acetylcholine; blue, GABA; yellow, glutamate; magenta, orphan GPCRs. *goa-1(n1134) I* was used as a control hyperactive egg-laying mutant. For panels A-C, n=100 eggs per strain. Statistical significance was determined with the Fisher’s exact test for the early-staged egg assay. The Wilson-Brown Method was used to determine the 95% C.I. for binomial data. p <0.05 (*) was considered significant. (B) Overexpression of the serotonin GPCR gene *ser-1* causes a strong hyperactive egg-laying defect that is reproduced in strains carrying an extrachromosomal (Ex.) *ser-1* transgene and three independent chromosomally-integrated *ser-1* transgenes. Control is WT. (C) One strain carrying an integrated transgene for the GABA GPCR *gbb-1* shows a strong hyperactive egg-laying defect that is not reproduced in a second chromosomal integrant or in a strain carrying an extrachromosomal (Ex.) *gbb-1* transgene. Controls are WT or a strain with fluorescent neurons carrying a *rab-3p::NLS::TagRFP* transgene. (D) Overexpression of the octopamine GPCR gene *ser-6* causes strong egg-laying defects as seen by the accumulation of unlaid eggs in the uterus in strains carrying four independent chromosomally-integrated *ser-6* transgenes and one extrachromosomal (Ex.) *ser-6* transgene. The strain carrying the integrated transgene *ser-6(oe)* #1 was outcrossed to wild type to remove the *rab-3p::NLS::TagRFP* transgene and re-assayed. Controls are WT or *rab-3p::NLS::TagRFP*; Brown: *ser-6* overexpressors. For this panel and panels E-I, statistical significance was tested using one-way ANOVA with a Tukey’s test for multiple comparisons. n ≥ 30 for each strain. p <0.05 (*) was considered significant. (E) Overexpression of the glutamate GPCR gene *mgl-1* causes strong egg-laying defects as seen by the accumulation of unlaid eggs in the uterus in both an extrachromosomal (Ex.) and chromosomally-integrated *mgl-1* transgenic strains. Controls are WT or *rab-3p::NLS::TagRFP*. (F) Overexpression of the glutamate GPCR gene *mgl-3* causes mild egg-laying defects as seen by the accumulation of unlaid eggs in the uterus in three extrachromosomal and one chromosomally-integrated *mgl-3* transgenic strains for MGL-3. Control is WT. (G) Overexpression of the acetylcholine GPCR gene *gar-1* reveals mild egg-laying defects as seen by the accumulation of unlaid eggs in the uterus in extrachromosomal (Ex.) and chromosomally-integrated *gar-1* transgenic strains. Control is WT. (H) Extrachromosomal (Ex.) overexpression of the octopamine GPCR gene *ser-3* causes strong egg-laying defects as seen by the accumulation of unlaid eggs in the uterus, but this phenotype is not reproduced in by a strain with a chromosomally-integrated *ser-3* transgene. Control is WT. (I) Overexpression of the tyramine GPCR gene *ser-2* causes mild egg-laying defects as seen by the accumulation of unlaid eggs in the uterus in extrachromosomal and chromosomally-integrated *ser-2* transgenic strains. Controls are WT and *rab-3p::NLS::TagRFP*.

Video S1: Methods Used to Score Expression of a GFP Reporter for the Octopamine Receptor OCTR-1 in Cells of the Egg-Laying System, Related to Figure 3

